# Fixel-wise differences in White Matter Fiber-bundle Cross-section identified in Children who Stutter

**DOI:** 10.64898/2026.10.29.762844

**Authors:** Hasini Weerathunge, Yanni Liu, Fiona Höbler, Mike Angstadt, Andrew Jahn, Ho Ming Chow, Soo-Eun Chang

## Abstract

Developmental stuttering has been repeatedly linked to decreased white matter integrity and structural connectivity differences in major white matter tracts. Prior diffusion-weighted MRI studies reporting such white matter structural differences involved voxel-averaged tensor measures (e.g., fractional anisotropy (FA)). A well-acknowledged limitation of this approach, however, has been that it cannot resolve distinct fiber populations within crossing-fiber regions. Fixel-based analysis addresses this limitation by quantifying individual fiber populations within a voxel and providing complementary measures of fiber density (FD), fiber-bundle cross-section (FC), and their combined measure (FDC).

Here we applied Fixel-based analysis to diffusion MRI data from 171 children aged 3–13 years, including 84 children who stutter and 87 typically developing controls, to test whether white-matter differences associated with developmental stuttering were expressed as differences in FD, FC, and FDC and whether these measures show group-specific age-related patterns. We evaluated this at both the whole-brain level and within a priori white-matter tracts.

FC, but not FD or FDC, showed significant Group and Group × Age effects, indicating that white-matter differences were primarily expressed in fiber-bundle macrostructure. Tract-of-interest analyses identified developmental differences in bilateral arcuate fasciculus FC that remained significant across tracts, with controls showing greater age-related increases in FC than children who stutter. Additional FC differences were observed across corticospinal and thalamocortical pathways, although these tract-wise effects were less robust to across-tract correction.

These findings suggest that white-matter differences in developmental stuttering are prominently expressed in fiber-bundle macrostructure rather than fiber density and highlight divergent development of bilateral arcuate pathways. FBA provides fiber-specific information that complements conventional tensor-based diffusion measures and may improve characterization of white-matter development in childhood stuttering.

**Significance Statement:** Developmental stuttering has repeatedly been associated with white-matter differences, but conventional diffusion measures cannot readily distinguish changes in individual fiber populations within regions containing crossing fibers. Using Fixel-based analysis in children who stutter, we show that these differences are primarily expressed in fiber-bundle cross-section rather than fiber density, implicating macrostructural differences in white-matter organization. The strongest tract-wise developmental effects were observed bilaterally in the arcuate fasciculus, a major pathway supporting auditory-motor integration, where children who stutter showed divergent age-related trajectories relative to typically developing children. These findings provide a fiber-specific perspective on the neurodevelopmental basis of stuttering and refine the interpretation of white-matter differences previously identified using conventional diffusion imaging.

## 1. Introduction

Developmental stuttering (hereafter referred to as stuttering) is a neurodevelopmental speech disorder that emerges in early childhood, affecting 5-7% of children and 1% of the general population (Gattie et al., 2025; Yairi & Ambrose, 2013). Stuttering can have a significant psychosocial impact throughout the lifespan and affect quality of life (Boyle & Cheyne, 2024; Craig et al., 2009; Johnson et al., 2024).

Over three decades of neurophysiological evidence have shed light on the potential neural underpinnings of stuttering (Chang et al., 2025; Etchell et al., 2018; Neef & Chang, 2024). Neuroimaging studies have shown both brain structure and function-based differences in children and adults who stutter when compared to age- and sex-matched participants (Neef et al., 2015). However, neurodevelopmental disorders are affected not only by initial deviations in neurophysiology but also by compensatory alterations in brain circuits throughout development (Chow et al., 2023; Chow & Chang, 2017a). Many adults who stutter exhibit functional or structural brain changes that may mask the initial core deficits associated with stuttering (Belyk et al., 2017; Loucks et al., 2011; Nil et al., 2000). Examining children closer to stuttering onset therefore provides an opportunity to characterize neurodevelopmental differences before decades of experience and adaptation have occurred (Chow et al., 2023; Chow & Chang, 2017a). White-matter architecture is particularly relevant in this context because it undergoes substantial maturation throughout childhood and provides the backbone for structural connectivity that supports developing speech networks. Prior diffusion MRI studies in CWS have identified atypical white-matter organization and developmental trajectories, motivating more detailed characterization of the fiber pathways and structural properties underlying these differences.

### 1.1 Neuroanatomical differences in developmental stuttering

Previous lines of research in brain structure in children who stutter are based on both T1-weighted structural MRI imaging, as well as diffusion-weighted imaging (diffusion MRI; Beal et al., 2013; Chang et al., 2008, 2015; Chow & Chang, 2017b; Matsuhashi et al., 2023). Structural MRI measures macrostructural features of the brain anatomy – the gray and white matter volume, cortical thickness, surface area, and gyral anatomy. Although detailed and provided in a high resolution (1 mm^3^), structural MRI provides relatively gross anatomical descriptors that are useful in detecting large-scale structural, volumetric, or cortical variations (Short et al., 2022; Tae et al., 2025). The white matter tracts constitute the major neural pathways that connect cortical and subcortical gray matter that allow rapid neural signaling throughout the brain. To understand and characterize neural circuits better, techniques that reveal the microstructure and integrity of these white matter tracts are needed.

Diffusion-weighted imaging measures the water diffusion along brain structures. Because water diffuses anisotropically within white matter – constrained by myelinated axonal membranes, water molecules move primarily parallel, not anti-parallel, to the length of the axon–diffusion MRI metrics are sensitive to axonal integrity and can enable both delineation of specific white matter tracts and the characterization of their microstructural and morphological properties, such as fiber density, orientation, and cross-section, which appear homogeneous on structural MRI (Baliyan et al., 2016; Bammer, 2003). Diffusion MRI offers a rich set of metrics reflecting tissue microstructure and are sensitive to early variations occurring in development, disease, and brain plasticity (Le Bihan & Johansen-Berg, 2012). Depending on the diffusion model applied — ranging from diffusion tensor-based approaches (DTI) to more advanced frameworks such as Fixel-based analysis (FBA) and Neurite Orientation Dispersion and Density Imaging (NODDI) — diffusion MRI can characterize white matter at both microstructural (fiber density, myelination) and macrostructural (tract cross-sectional morphology of tracts, connectivity) levels, the latter being particularly important for understanding how tract morphology relates to capacity for signal transmission across development and disease. However, conventional diffusion MRI models such as DTI can be limited in two key aspects: these methods are susceptible to partial volume effects from gray matter and CSF and may provide less stable estimates in regions of complex fiber architecture, such as areas with crossing fibers (J. -D. Tournier et al., 2002).

To date, relatively few neuroimaging studies have focused on examining childhood stuttering (Chang et al., 2008; Chang & Zhu, 2013; Choo et al., 2012). The DTI studies published to date have identified reduced FA across multiple tracts in individuals who stutter, spanning both pediatric and adult samples, though the precise location of these reductions has been inconsistent across studies (Neef et al., 2015) – likely reflecting methodological heterogeneity in participants’ age ranges, tract definition approaches, and the voxel-averaged limitations of FA itself. Reduced FA has been reported along the left superior longitudinal fasciculus (SLF) and arcuate fasciculus (AF) underlying the inferior frontal gyrus (Kell et al., 2009, adults; Chang et al., 2015, children), premotor cortex (Chang et al., 2008, children; Watkins et al., 2008, adolescents/adults; Connally et al., 2014, mixed-age), Rolandic operculum (Sommer et al., 2002, adults; Chang et al., 2008, children), and inferior parietal region (Chang et al., 2008, children; Watkins et al., 2008, adolescents/adults). FA reductions have also been noted along white matter tracts connecting cortical and subcortical regions, such as the CST (Cai et al., 2014, adults) and the CC (Cykowski et al., 2010, adults; Chang & Zhu, 2013, children; Chang et al., 2015, children). A meta-analysis by Neef and colleagues spanning this largely adult/mixed-age literature, reported convergence of FA reductions associated with developmental stuttering in the AF and the body of the CC (Neef et al., 2015).

More recently, Chow and Chang conducted a longitudinal DTI study comprising 35 CWS and 43 CNS, ranging in age from 3 – 10 years, aimed at identifying white matter microstructural differences occurring in children with persistent stuttering and to examine structural changes associated with recovery of childhood stuttering (Chow & Chang, 2017b). Reduced FA was found in the left AF, underlying the inferior parietal and posterior temporal areas, and the mid-body of the CC among CWS as a group relative to controls. Moreover, the authors reported a reduced FA growth rate in the same regions among children with persistent stuttering, which was not observed among those who recovered from stuttering. Collectively, these results highlight several white matter tracts relevant to the neural bases of childhood stuttering: frontal-motor to auditory connectivity subserving movement planning, execution, and sensorimotor integration; cortico-subcortical pathways between SMA and putamen supporting motor timing and sequence coordination; and callosal fibers enabling interhemispheric communication. The callosal anomalies found in particular suggest interhemispheric reorganization, consistent with observed differences in right-hemispheric gray matter morphology and alterations in the right SLF (Cai et al., 2014; Kronfeld-Duenias et al., 2016).

### 1.2 Advances in diffusion MRI analysis: Fixel-based analysis (FBA)

Although diffusion MRI provides a range of non-invasive and in vivo metrics that are sensitive to white matter microstructure, the cellular bases of these metrics remain incompletely understood. In particular, the relative contributions of axonal membranes, myelin sheaths, extracellular matrix, and other tissue compartments to the diffusion signal are not fully elucidated. These limitations are especially relevant for DTI, where several methodological caveats can complicate interpretation, particularly in regions of complex fiber geometry. Because DTI cannot resolve multiple fiber populations within a voxel, it does not accurately represent crossing-fiber configurations at a sub-voxel level (Dhollander et al., 2021a). In voxel-averaged analyses, this complexity is instead summarized by a single FA value per voxel, which can produce misleading FA estimates in regions containing crossing fibers. Moreover, DTI metrics are inherently non-specific to axonal properties, because the diffusion signal reflects contributions from both intra-axonal and extra-axonal compartments, including glial cells and extracellular space. In addition to FA reductions, Chow and colleagues (2017) identified regions showing increased FA and higher FA growth rates in both persistent and recovered CWS relative to controls; these regions were adjacent to areas showing reduced FA or FA growth rates (Chow & Chang, 2017a). Although several neurophysiological interpretations of these findings are possible, the observed FA increases may also partly reflect methodological artifact, because apparent increases in FA within one tract can arise from reduced FA in another tract at crossing-fiber junctions. The tract tracing may either seize at a crossing fiber junction or erroneously reduce FA values depending on how these junctions are handled (Jones & Cercignani, 2010).

Alternative multi-compartmental models, such as neurite orientation dispersion and density imaging (NODDI; Zhang et al., 2012), offer greater microstructural specificity by separately modeling intra-neurite and extra-neurite compartments. These models yield metrics such as neurite density index (NDI) and orientation dispersion index (ODI), which are more directly related to axonal and dendritic morphology than FA. However, NODDI remains a voxel-level model that typically represents neurite orientation as a single orientation distribution within each voxel. As a result, it cannot resolve the properties of distinct fiber populations in regions where multiple bundles intersect at different orientations (A. W. Anderson, 2005; Zhang et al., 2012; Dhollander et al., 2021b). Accordingly, methods that can resolve individual fiber populations within a voxel are needed to characterize white matter microstructure in a fiber-specific manner and to distinguish genuine microstructural differences from crossing-fiber and partial volume effects.

To address these challenges, a new fiber-specific analysis pipeline has been proposed (D. A. Raffelt et al., 2015, 2017) where each individual **<u>fi</u>**ber population within a vo**<u>xel</u>** (termed as **“Fixel”**) is resolved, allowing fiber-specific metrics to quantify white matter properties (Dhollander et al., 2021b). In FBA, white matter fiber orientation densities (FODs) are computed via constrained spherical deconvolution (CSD; J.-D. Tournier et al., 2008; Jeurissen et al., 2014; Dhollander & Connelly, 2016). A Fixel-wise apparent fiber density, more broadly known as fiber density (FD; D. Raffelt et al., 2012), can be derived from the FODs. FD is approximately proportional to total intra-axonal volume and is a measure of fiber-specific white matter microstructure (Genc et al., 2020; D. Raffelt et al., 2012). Macroscopic differences in fiber-bundle cross section (FC; D. A. Raffelt et al., 2017) can also be obtained via FBA at a Fixel-wise level. This is obtained by warping individual participant data to a common template space, resulting in an equivalent to traditional tensor-based morphometry (TBM; Ashburner & Friston, 2000), but at a Fixel-wise level. Fixel-wise FD and FC measures are also multiplied to form a combined fiber density and cross-section measure (FDC; D. A. Raffelt et al., 2017), which jointly captures both microstructural (axonal density) and macrostructural (tract morphology) contributors to white matter differences. FDC is therefore particularly sensitive to overall differences in the white matter fiber pathway capacity — reflecting the total number of axons coursing through a given tract cross-section – and is described to be broadly analogous to voxel-based morphometry (VBM; Ashburner & Friston, 2000).

Overall, Fixel-based metrics provide fiber-population-specific estimates of white matter micro- and macrostructure (see Figure 1). This feature is particularly important in regions of complex fiber architecture, where voxel-averaged DTI measures may conflate signals from multiple fiber populations. By resolving distinct fiber populations within a voxel, FBA can help clarify whether previously reported white matter differences in CWS reflect genuine fiber-specific alterations or partial volume effects related to crossing-fiber geometry. Because FA reductions in CWS have been reported primarily in tracts such as AF, SLF, CST, and CC, which are tracts that traverse regions of complex fiber architecture near areas such as the IFG, it remains uncertain whether these findings reflect true fiber-population-specific white matter alterations or partial volume effects arising from crossing fibers. The FBA approach addresses this limitation by separately estimating the properties of individual fiber populations within a voxel, providing a more specific framework for re-evaluating white matter differences in CWS relative to controls.

**Figure 1:**
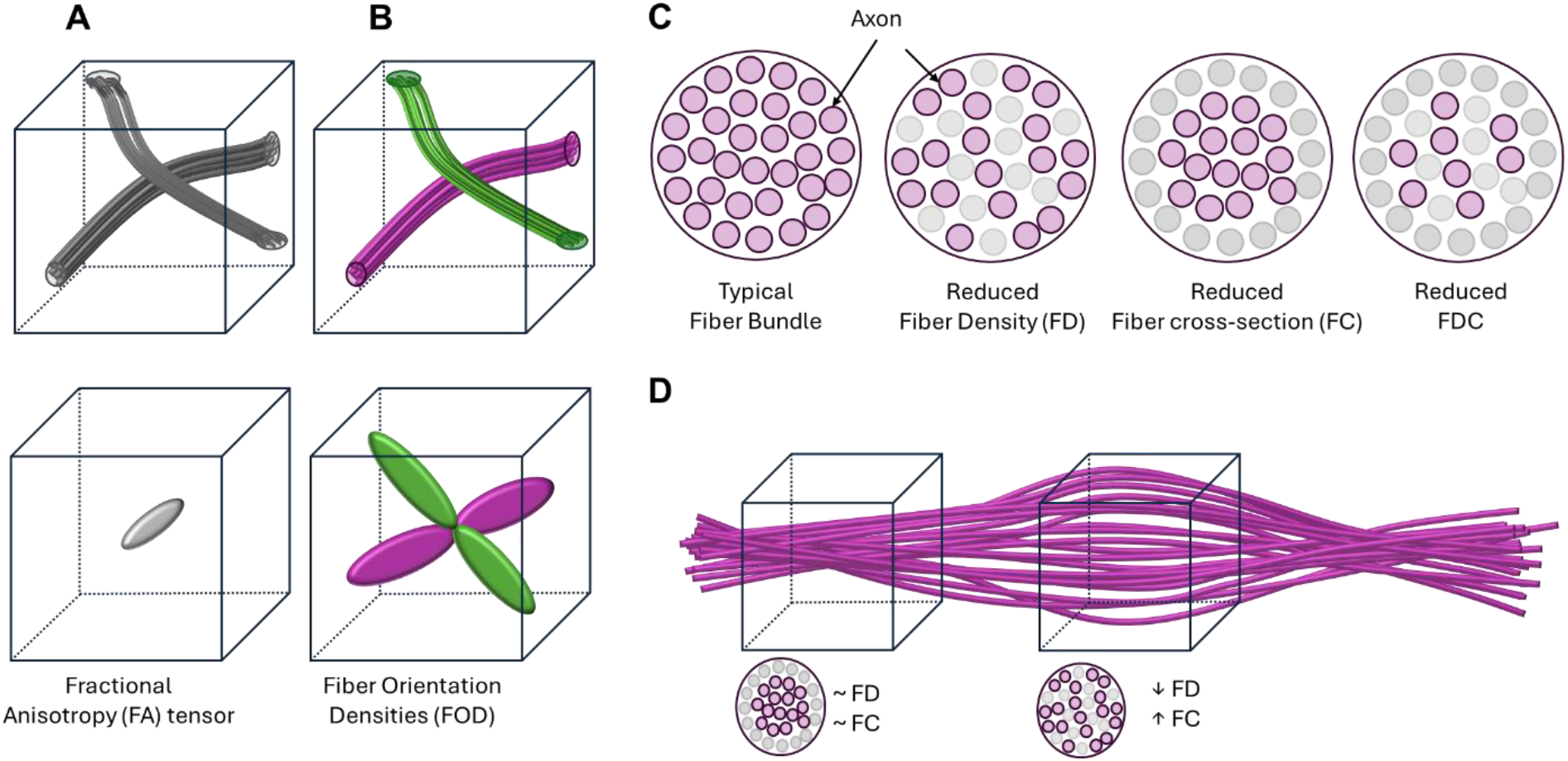
Conceptual illustration of Fixel-based measures and comparison with traditional voxel-averaged metrics. **A.** Conventional voxel-averaged tensor metrics, such as fractional anisotropy (FA), average diffusion signals across all fiber populations within a voxel, which can be obscured by factors such as crossing fibers. **B.** Fixel-based analysis separates distinct fiber populations (“Fixels”) within each voxel (magenta and green). **C.** Fiber density (FD) captures normalized intra-axonal fiber volume, while fiber bundle cross-section (FC) reflects macroscopic bundle morphology derived from nonlinear registration to a population template. Their combined metric, fiber density and cross-section (FDC), summarizes the overall white matter fiber pathway capacity. Collectively, FBA distinguishes microstructural versus macrostructural white matter alterations. **D.** Variations of microstructural and macrostructural properties captured via Fixel-based measurements are illustrated via differences in fiber density and cross-section across the multiple voxels.

Because FBA separately estimates fiber density (FD) and fiber cross-section (FC), it can help disentangle microstructural differences within fiber populations from macrostructural differences in tract morphology. These metrics therefore provide a more specific context for interpreting FA reductions previously reported in CWS, which may reflect altered fiber density (myelin-related microstructural alterations), altered tract morphology (macrostructural reorganization), or both. In neurodevelopmental disorders, white matter differences may reflect macrostructural alterations in fiber organization, including differences in tract morphology, cross-section, or trajectory, rather than, or in addition to, microstructural changes such as altered myelination (Kelly et al., 2020, 2020; Kirkovski et al., 2020). These microstructural differences may emerge from atypical developmental processes, including deviations in fiber pruning, fiber bundle consolidation, or tract trajectory refinement (Lebel & Beaulieu, 2011). Accordingly, FA reductions may not necessarily indicate reduced myelination, but instead reflect broader differences in white matter organization that influence diffusion anisotropy (Jones & Cercignani, 2010; D. A. Raffelt et al., 2017). In contrast to FA, which cannot provide sub-voxel level fiber resolution or the feasibility to separate micro- vs. macro-structural changes in white matter, Fixel-based measures of fiber density and cross-section allow separation of micro- vs. macro-level changes in white matter fiber architecture to provide insights into these components of white matter developmental trajectories. So far, there have been no studies exploring fixel-based measures in adults or children who stutter. The more traditional DTI-based tract-wise analysis methods also have limited tract-specific resolution. More recent methods, such as TractSeg (Wasserthal et al., 2018), leverage deep learning to delineate major white matter pathways with greater specificity, using tract templates derived from the JHU white matter atlas (“MRI Atlas of Human White Matter,” 2006). The present study combines TractSeg-based tract delineation with FBA to provide the first fiber-specific, tract-localized characterization of micro- and macrostructural white matter differences in CWS relative to typically developing peers.

### 1.3 The current investigation

In this study, we aimed to characterize fiber-specific alterations in white matter architecture in children who stutter (CWS) using Fixel-based analysis (FBA). While previous DTI studies have identified differences in white matter integrity in CWS, tensor-derived measures such as fractional anisotropy (FA) cannot distinguish between different fiber populations within crossing-fiber regions or determine whether these differences primarily reflect fiber-specific microscopic or macroscopic morphological properties. We therefore examined, for the first time in this clinical population, fiber density (FD; reflecting fiber-specific microstructure), fiber-bundle cross-section (FC; reflecting macroscopic fiber-bundle morphology), and their combined measure, fiber density and cross-section (FDC).

We addressed two primary research questions. First, we investigated **whether fiber-specific white matter architecture differs between children who stutter (CWS) and typically developing controls**. Based on previous DTI-based reduced FA findings, we hypothesized that the CWS would exhibit alterations in the white matter morphology within pathways implicated in speech and motor function, particularly the arcuate fasciculus (AF)/superior longitudinal fasciculus (SLF), which are critical for auditory-motor integration and speech planning; the corticospinal tract (CST), indicating deficits in motor coordination and timing; and the corpus callosum (CC), inferring deficits in interhemispheric communication. By separately examining FD and FC, we sought to determine whether previously reported diffusion abnormalities were predominantly associated with differences in fiber-specific microstructure reflected by FD, macroscopic fiber-bundle morphology reflected by FC, or a combination of these properties, as reflected by FDC. Second, we investigated **whether associations between age and Fixel-based white matter measures differed between groups.** Based on previous findings of reduced age-related increases in white matter integrity in children with persistent stuttering relative to typically developing controls, we hypothesized weaker positive age associations in CWS, particularly within the AF and CC.

Finally, we examined **associations between Fixel-based measures and stuttering severity in CWS**. Based on previous reports linking lower white matter integrity (FA) in speech-and motor-related pathways with greater stuttering severity, we hypothesized that lower Fixel-based measures would be associated with greater stuttering severity in CWS (left IFG, left precentral gyrus/M1, and left supramarginal gyrus; Chang et al., 2015; anterior CC; Civier et al., 2015).

## 2. Materials and Methods

### 2.1 Participants

All participants were native English speakers with no history of neurological or psychiatric disorders. Details on behavioral measurement procedures and a list of assessments can be found in previous studies (Chang et al., 2015; Chang & Zhu, 2013; Chow & Chang, 2017a). All procedures used in the study were approved by the Michigan State University Institutional Review Board. The CWS and CNS were age- and sex-matched with no difference across groups in socioeconomic status (Hollingshead, 1975) or verbal Intelligence quotient (IQ). See Table 1 for participant demographic information.

**Table 1.** Participant Demographic Information.

|  | Control (n = 87; 44 males) |  | Stuttering (n = 84; 51 males) |  | t-value | p (two-tailed) |
| --- | --- | --- | --- | --- | --- | --- |
|  | Mean (SD) | Range | Mean (SD) | Range |  |  |
| Age at MRI (months) | 66.93<br>(21.64) | 37 – 129 | 67.92<br>(23.75) | 37 – 158 | – 0.283 | .777 |
| SES | 50.41<br>(10.48) | 18 – 66 | 49.85<br>(10.85) | 18 – 66 | 0.341 | .733 |
| VIQ | 114.63<br>(14.01) | 87 – 153 | 108.68<br>(13.54) | 77 – 137 | 2.826 | **.005\*** |
| TIV (mm<sup>3</sup>) | 1385.05<br>(115.20) | 1090.33 – 1750.33 | 1398.17<br>(135.49) | 1152.64 – 1810.17 | – 0.682 | .497 |
| Mean FD (mm) | 0.64<br>(0.29) | 0.31 – 1.72 | 0.82<br>(0.41) | 0.27 – 2.37 | – 3.34 | **.001\*** |
| SSI-4 at initial visit | N/A | N/A | 19.14<br>(7.05) | 2 – 48 | N/A | N/A |

SES: Socioeconomic status; VIQ: Verbal intelligence quotient; TIV: Total Intracranial Volume; FD: Framewise Displacement; SSI-4: Stuttering severity instrument Edition 4.
**Note.** The stuttering group comprised 62 children with persistent stuttering and 22 children who recovered.

Stuttering severity was assessed utilizing a series of speech samples collected during spontaneous speech tasks (conversational tasks and storytelling conducted with a parent and a certified speech-language pathologist). These speech samples were video-recorded and analyzed offline according to the procedures listed in the Stuttering Severity Instrument Edition 4 (SSI-4; Riley, 2009). All children who stutter (n = 84) were diagnosed with stuttering during their initial visit but later categorized as persistent (n = 62) or recovered (n = 22) based on scored stuttering severity and frequency based on speech samples, parent report, and clinician assessments during subsequent visits. Specifically, a child who stutters was considered as persistent if their %SLD was 3 or higher and composite SSI-4 score was higher than 10 at two follow-up visits. A child who stutters was considered recovered if the percent stuttering-like disfluencies (%SLD) was lower than 3 and composite SSI-4 score was below 10 (considered “very mild” in SSI-4 stuttering severity classification) at two consecutive follow-up visits. The classification of persistence or recovery was based not only on speech assessment but also considered clinician impression and parental reports. These classifications corroborated the prior studies’ criteria for persistent vs. recovery categorization (Ambrose & Yairi, 1999).

### 2.2 MRI Data Acquisition

Diffusion MRI and structural MRI (T1-weighted) data were acquired across two cohorts on GE 3T Signa® HDx MR scanners (GE Healthcare, Chicago, IL, USA) with an 8-channel head coil. Cohort 1 comprised 82 participants (40 controls and 42 children who stutter; age range, 37–158 months), whereas Cohort 2 comprised 89 participants (47 controls and 42 children who stutter; age range, 37–124 months; See Supplemental Table 1). Because age distributions and acquisition parameters differed between acquisition cohorts, acquisition cohort was included as a nuisance covariate in all group-level statistical models.

For cohort 1, diffusion-weighted images were acquired with a dual spin-echo echo-planar imaging sequence with the following parameters. 48 contiguous 2.4 mm axial slices in an interleaved order, field of view = 22 x 22 cm, matrix size = 128 x 128, number of excitations = 2, echo time = 77.5 ms, repetition time = 13.7 s, 25 diffusion-weighted volumes (one per gradient direction) with b = 1,000 s/mm^2^, and one volume with b = 0. For cohort 2, diffusion-weighted images were acquired with a dual spin-echo echo-planar imaging sequence with the following parameters. 48 contiguous 2.4 mm axial slices in an interleaved order, field of view = 22 x 22 cm, matrix size = 128 x 128, number of excitations = 1, echo time = 84.8 ms, repetition time = 12.8 s, 40 diffusion-weighted volumes (one per gradient direction) with b = 1,000 s/mm^2^, and 4 volumes with b = 0.

During each session, 180 T1-weighted 1-mm^3^ isotropic volumetric inversion recovery fast spoiled gradient-recalled images, with CSF suppressed, were obtained to cover the whole brain with the following parameters: time of echo = 3.8 ms, time of repetition of acquisition = 8.6 ms, time of inversion = 831 ms, repetition time of inversion = 2,332 ms, flip angle = 8°, and receiver bandwidth = ±20.8 kHz.

For all scans across the cohorts, one staff member sat in the scanner room next to the child at all times to monitor the child’s comfort and to ensure cooperation during scanning. During the acquisition of volumetric T1-weighted and DTI scans, the children viewed a movie to help them stay still.

To ensure adequate diffusion MRI data quality, all diffusion-weighted scans underwent quality control procedures prior to inclusion in the final analysis. Scans with excessive motion-related corruption, signal dropout, or other visible image artifacts were excluded from further analysis. As an additional quantitative quality-control assessment, slice-level signal outliers identified during Functional Magnetic Resonance Imaging of the Brain Software Library (FSL) *eddy* current and motion correction were examined using the corresponding *eddy* outlier map. *Eddy* uses a model-based approach to predict the expected diffusion signal from the acquired diffusion measurements and identifies slices showing unusually large deviations between the observed and predicted signal, which can reflect motion-related signal dropout and other signal corruption. Using the default eddy outlier criterion, slices with residuals exceeding the expected value by more than four standard deviations were classified as outliers. For each diffusion volume, the proportion of slices classified by *eddy* as outliers was calculated, and volumes in which more than 20% of slices were identified as outliers were considered susceptible to substantial signal corruption. Consistent with the scan-level criterion used in prior work for a subset of the current dataset (Chow & Chang, 2017a), scans containing more than 10 such volumes were considered for exclusion from further analysis.

Following quality control, a total of 171 high-quality DTI scans from 84 CWS (male = 51; mean age (months) = 67.92 (23.75)) and 87 CNS (male = 44; mean age (months) = 66.93 (21.64)) between 3 and 13 years of age were retrospectively analyzed for this cross-sectional study. One scan was included per participant. The earliest available scan satisfying diffusion MRI quality-control criteria was selected; when the initial scan did not meet these criteria, a subsequent longitudinal scan was used.

### 2.3 Whole-brain Fixel-Based Analysis

Data were preprocessed in the participant’s native space, including denoising, motion correction, susceptibility distortion correction, and bias field correction using MRtrix3 (Version RC3; J.-D. Tournier et al., 2012) according to the FBA pipeline provided by MRtrix3 documentation. Corrections for subject motion and eddy-current-induced distortions were specifically performed using FSL *eddy* within the diffusion preprocessing workflow for MRtrix3 software. Residual head motion within each participant’s diffusion acquisition was quantified from the six rigid-body motion parameters estimated by *eddy*. These estimates were summarized at the participant level to generate a motion measure to account for between-participant differences in head motion in subsequent statical analyses.

Following preprocessing, diffusion MRI data were processed using MRtrix3 (Version RC3; J.-D. Tournier et al., 2012) according to the Fixel-based analysis (FBA) pipeline. Tissue-specific response functions for white matter (WM), grey matter (GM), and cerebrospinal fluid (CSF) were first estimated separately for each participant in acquisition cohort 2 using the unsupervised response function estimation method (Dhollander et al., 2016, 2019). Cohort 2 was selected for response-function estimation because it provided greater diffusion angular sampling than cohort 1 (40 versus 25 diffusion-weighted directions). These participant-specific response functions were then averaged to obtain representative WM, GM, and CSF response functions. As cohort 2 included participants from both the CWS and control groups, the resulting average response functions were independent of diagnostic group and were applied uniformly to participants across both groups and acquisition cohorts. These response functions were subsequently used to estimate WM fiber orientation distributions (FODs), together with GM-like and CSF-like signal compartments in all voxels (Dhollander & Connelly, 2016), from the preprocessed single-shell diffusion data using Single-Shell 3-Tissue Constrained Spherical Deconvolution (SS3T-CSD), implemented in MRtrix2Tissue (https://3Tissue.github.io), a fork of MRtrix3 (J.-D. Tournier et al., 2019). The resulting tissue compartments were subsequently subjected to joint bias-field correction and multi-tissue intensity normalization.

A study-specific population template for WM-FODs was generated using all 171 participants for group-wise comparison. Fixel-based measures of fiber density (FD), fiber cross-section (FC), and combined fiber density and cross-section (FDC) were computed following the MRtrix3 FBA pipeline. FD was calculated for each Fixel from the integral of the corresponding FOD lobe. FC was derived from the formation fields obtained during nonlinear registration of each participant’s FOD image to the population FOD template. FC values were log-transformed (logFC) pior to statistical analysis. FDC was calculated as the product of FD and the untransformed FC measure (FD x FC). Participant Fixel data was subsequently transformed into population template space for group-level statistical analysis (see the FBA pipeline documentation provided by MRtrix software for further details). Group differences were analyzed Fixel-wise using General Linear Models (GLMs). The full analysis workflow is outlined in Supplemental Figure 1. JHU white matter region and tract atlases provided via FSL were used to overlay significant Fixel regions with regions of interest (ROIs).

### 2.4 Tract of Interest (TOI) Analysis

Standard whole-brain Fixel-based group comparison provides the greatest spatial specificity; thus, we applied tract-specific analyses to comprehensively investigate how specific tracts of interest are implicated and in what way. Moreover, by constraining comparisons to only those Fixels that belong to *a priori* defined tracts, we can reduce the number of comparisons considerably and increase the statistical power. All analyses were conducted similarly to whole-brain Fixel-wise analyses except that each Fixel metric calculation was restricted to the specific tract-of-interest using Fixel masks.

White matter tracts were segmented in population-template space using TractSeg (Wasserthal et al., 2018). The selection of the segmented tract set was guided by previous DTI studies in stuttering, and included association tracts (Intrahemispheric Connectors: Arcuate Fasciculus [AF], Inferior Longitudinal Fasciculus [ILF], and Superior Longitudinal Fasciculus I – III [SLF I – III]), projection tracts (Cortico-Subcortical Connectors: Anterior Thalamic Radiation [ATR], Superior Thalamic Radiation [STR], Corticospinal Tract [CST]), and commissural fibers of the Corpus Callosum (CC). See Table 2 for more details. These TractSeg-derived segmentations were used to define the tract masks for the tract-of-interest (TOI) FBA analyses described below and to anatomically characterize the distribution of significant Fixels identified in the whole-brain analyses.

**Table 2.** A priori tract selection and rationale for primary and additional tract-of-interest analyses.

| **Tract** | **Anatomical Role** | **Rationale in Stuttering** | **Supporting Literature** |
| --- | --- | --- | --- |
| **Corpus Callosum (CC)** | Interhemispheric cortical connectivity | Reduced FA and atypical development in callosal motor regions | Chang et al., 2015; Chow & Chang, 2017 |
| **Corticospinal Tract (CST)** | Descending motor pathway | Altered integrity, particularly in left motor pathways | Kronfeld-Duenias et al., 2016 |
| **Arcuate Fasciculus (AF)** | Dorsal frontotemporal language pathway | Reduced FA and atypical developmental trajectories; auditory–motor integration | Chang et al., 2015; Chow et al., 2018 |
| **Superior Longitudinal Fasciculus (SLF) III** | Ventral frontoparietal pathway | Sensorimotor integration relevant to speech | Makris et al., 2005; Cai et al., 2014 |
| **Anterior Thalamic Radiation (ATR)** | Frontal thalamocortical pathway | Atypical development; relevant to CBGT circuitry | Chow & Chang, 2017; Chang & Guenther, 2019 |
| **Superior Thalamic Radiation (STR)** | Sensorimotor thalamocortical pathway | Atypical development; relevant to speech-motor circuitry | Chow & Chang, 2017; Chang & Guenther, 2019 |
| *Note: Eleven prespecified tracts constituted the primary tract-of-interest (TOI) analysis and the family used for BH-FDR correction. Six additional tracts examined outside the primary TOI set are indicated by grey shading.* |  |  |  |
| **Superior Longitudinal Fasciculus (SLF) I** | Dorsal frontoparietal pathway | Higher-order motor and attentional control | Makris et al., 2005; Palacios et al., 2021 |
| **Superior Longitudinal Fasciculus (SLF) II** | Prefrontal–parietal pathway | Attentional integration; implicated in stuttering | Makris et al., 2005; Neef et al., 2018 |
| **Inferior Longitudinal Fasciculus (ILF)** | Ventral occipitotemporal pathway | Atypical developmental trajectory reported in childhood stuttering | Chow & Chang, 2017 |

TractSeg requires three principal fiber orientations per voxel as input. Accordingly, fiber-orientation peaks were extracted from the study-specific WM-FOD population template generated during the preceding FBA processing steps (Jeurissen et al., 2014; J.-D. Tournier et al., 2008). A maximum of three peaks per voxel were extracted; when fewer than three fiber populations were present within a voxel (e.g., CC fibers), the remaining peak components were set to zero. Because peak extraction was performed directly from the study-specific WM-FOD template, the resulting TractSeg segmentations were generated in the same population template space used for the FBA.

Tract selection was guided by prior neuroimaging evidence and the anatomical relevance of specific white-matter pathways to speech motor control and developmental stuttering. A hypothesis-driven subset comprising bilateral AF, ATR, STR, CST, and SLF III and the CC (11 masks total) was used for formal TOI statistical inference. These pathways were prioritized based on their direct relevance to auditory–motor, speech-motor, corticospinal, interhemispheric, and thalamocortical circuitry implicated in developmental stuttering. ILF, SLF I, and SLF II were retained for anatomical characterization but were not included in the formal TOI multiple-comparison family. The anatomical and neurobiological rationale for the examined pathways is summarized in Table 2.

Anatomical localization of Fixel-wise effects was primarily obtained by tract-level interpretation of their spatial distribution within the template space. To aid descriptive reporting, voxel-wise overlap with the Johns Hopkins University (JHU) white matter tractography atlas and the JHU International Consortium for Brain Mapping (ICBM) Diffusion Tensor Imaging (DTI) 81 White matter labels atlas was additionally calculated (Hua et al., 2008; Mori et al., 2009; Wakana et al., 2007), although interpretation was guided primarily by tract-level correspondence of significant Fixels.

### 2.5 Statistical Analysis

#### 2.5.1 Whole-Brain Fixel-Based Analysis

Fixel-based measures of fiber density (FD, log-transformed fiber cross-section (logFC), and combined fiber density and cross-section (FDC) were analyzed via whole-Brain Fixel-wise General Linear Models (GLMs). GLMs were applied at each Fixel to identify the main effects of group (CWS vs. Control) and group x age interaction. Continuous covariates included age, total intracranial volume (TIV), socioeconomic status (SES), verbal IQ composite score (VIQ), and the participant-level head-motion measure described above; all continuous covariates were mean-centered prior to inclusion in the models. Sex and diffusion MRI acquisition cohort were included as coded categorical covariates. The group x age interaction term was constructed using mean-centered age.

All Fixel-wise analyses were conducted using the FBA pipeline (Dhollander et al., 2021b). Connectivity-based Fixel enhancement (CFE) was used to generate connectivity-based smoothing and statistical inference at the whole-brain level (CFE; D. A. Raffelt et al., 2015). CFE uses probabilistic tractography to identify structurally connected Fixels that are likely to belong to the same fiber tracts. This probabilistic connectivity information is then used for tract-specific smoothing and enhancement of the statistical map. Null distributions were generated using 5000 permutations of the design matrix within the GLM framework to estimate the distribution of the Fixel-wise test statistics under the null hypothesis. Permutation-based inference was combined with connectivity-based Fixel enhancement to exploit anatomical connectivity between Fixels and improve statistical power. Family-wise error (FWE)-correction was applied to generate corrected p-values (*p_FWE_* < .05). Based on prior neuroimaging studies in adults and children who stutter (limited clinical datasets), an exploratory uncorrected threshold of *p*_unc_ < .001 was additionally used to identify candidate Fixels for descriptive cluster-level characterization. For each GLM, directional contrasts were tested separately for CWS > CNS and CWS<CNS to identify both increases and decreases in FC, FD, and FDC associated with group and group × age interaction effects.

Following the whole-brain analysis, clusters of significant Fixels (voxel volume > 10 mm³, *p_unc_* < .001) were identified using the 3dClust function of the Analysis of Functional Neuroimages toolbox (AFNI; Cox, 1996) to characterize the anatomical extent and location of effects. No additional statistical tests or effect sizes are reported for this step, as any values derived from data used to define the cluster would be non-independent and potentially inflated (Kriegeskorte et al., 2010).

#### 2.5.2 Tract of Interest (TOI) Analysis

Tract of Interest (TOI) GLMs were conducted across five tracts of interest defined as per the JHU white matter atlas: Arcuate Fasciculus, Anterior Thalamic Radiation, Superior Thalamic Radiation, Corticospinal Tract, Corpus Callosum, and Superior Longitudinal Fasciculus III (five bilateral, one unilateral [Corpus Callosum]; 11 tracts total).

To account for multiple comparisons across the 11 a priori tract-based analyses, a two-stage correction procedure was used. Connectivity-based Fixel enhancement was performed separately within each tract mask using 5,000 permutations, yielding Fixel-wise family-wise error (FWE)-corrected *p* values within each tract. For each directional contrast, the minimum FWE-corrected *p* value within each tract was extracted and the resulting 11 tract-level *p* values were subjected to Benjamini–Hochberg false discovery rate (BH-FDR) correction at *q* < .05. BH-FDR correction was performed separately for the positive and negative Group contrasts and for the positive and negative Group × Age interaction contrasts. To characterize the spatial extent of effects within each tract, the number and percentage of Fixels reaching within-tract FWE-corrected *p* < .05 were also calculated. These extent measures were descriptive and were not used as inputs to the BH-FDR correction.

### 2.6 Exploratory Analyses

#### 2.6.1 Relationship with Stuttering Severity

Within the group of CWS, we examined whether Fixel-based measures correlated with stuttering severity measured with SSI-4 (Riley, 2009). Mean Fixel values across significant Fixel clusters were plotted against SSI-4 scores for persistent and recovered children who stutter separately. Scatter plots indicated the direction and magnitude of association. Similarly, scatter plots were generated for tract-wise analyses.

To further explore heterogeneity within the group of children who stutter, we also conducted a secondary whole-Brain Fixel-Wise GLM, treating typically developing children, persistent, and recovered subgroups of children who stutter as separate subgroups. No significant Fixels survived corrections for multiple comparisons at the whole-brain level.

#### 2.6.2 DTI-based fractional Anisotropy (FA) measure

To assess the consistency of Fixel-Based findings with traditional diffusion tensor imaging (DTI) metrics, we conducted a parallel analysis using Fractional Anisotropy (FA) derived from Tract-Based Spatial Statistics (TBSS) on the same dataset. Group differences and Group x Age interactions were modeled using a general linear model (GLM) that included the same covariates as the FBA. Statistical inference was performed using FSL’s randomize with 5,000 permutations and threshold-free cluster enhancement.

## 3. Results

### 3.1 Whole-brain Fixel-wise Analyses

For whole-brain Fixel-based analyses, only Fiber Cross-Section (FC) showed significant effects of group and group x age interactions at *p_unc_* <.001. Neither Fiber Density (FD) nor Fiber Density and Cross-Section (FDC) showed significant effects. Among the FC results, only the Group x Age interaction (CWS < Controls) survived FWE corrections at *p_FWE_* <.05. Group effects did not survive Family-wise error correction, but uncorrected p-value maps (at *p_unc_* <.001 and *p_unc_* <.005) are shown for illustrative purposes (Group effects: Supplemental Figure 2, Group x Age Interaction: Supplemental Figure 4). Anatomical Localization of the whole-brain results using JHU and ICBM white matter atlases showed that group differences showing CWS>controls for FC were localized to right anterior thalamic radiation, right inferior fronto-occipital fasciculus, and to the right anterior corona radiata.

Clusters exceeding a voxel volume of > 10 voxels at *p_unc_* <.001 were included in subsequent cluster-wise analyses (see Figure 2 for Group x Age interaction [CWS < controls: effects and clusters]. Seven significant clusters resulted from the Group x Age interactions, five of which overlapped with tracts of interest. Among these, two overlapped with multiple tracts of interest (corresponding to crossing-fiber regions), and three with a single tract of interest (Supplemental Table 2). Cluster 1 overlapped with Arcuate Fasciculus (left; 31.49%), Inferior Longitudinal Fasciculus (left; 7.79%), Corpus Callosum (6.79%), and Superior Longitudinal Fasciculus II (left; 0.03%) tracts. Cluster 2 overlapped with Arcuate Fasciculus (right; 42.45%), Corpus Callosum (5.91%), and Inferior Longitudinal Fasciculus (right; 3.60%). Clusters 3,4, and 5 overlapped with Inferior Longitudinal Fasciculus (left; 0.26%), Corpus Callosum (3.66%), and Anterior Thalamic Radiation (left; 8.40%), respectively.

**Figure 2.**
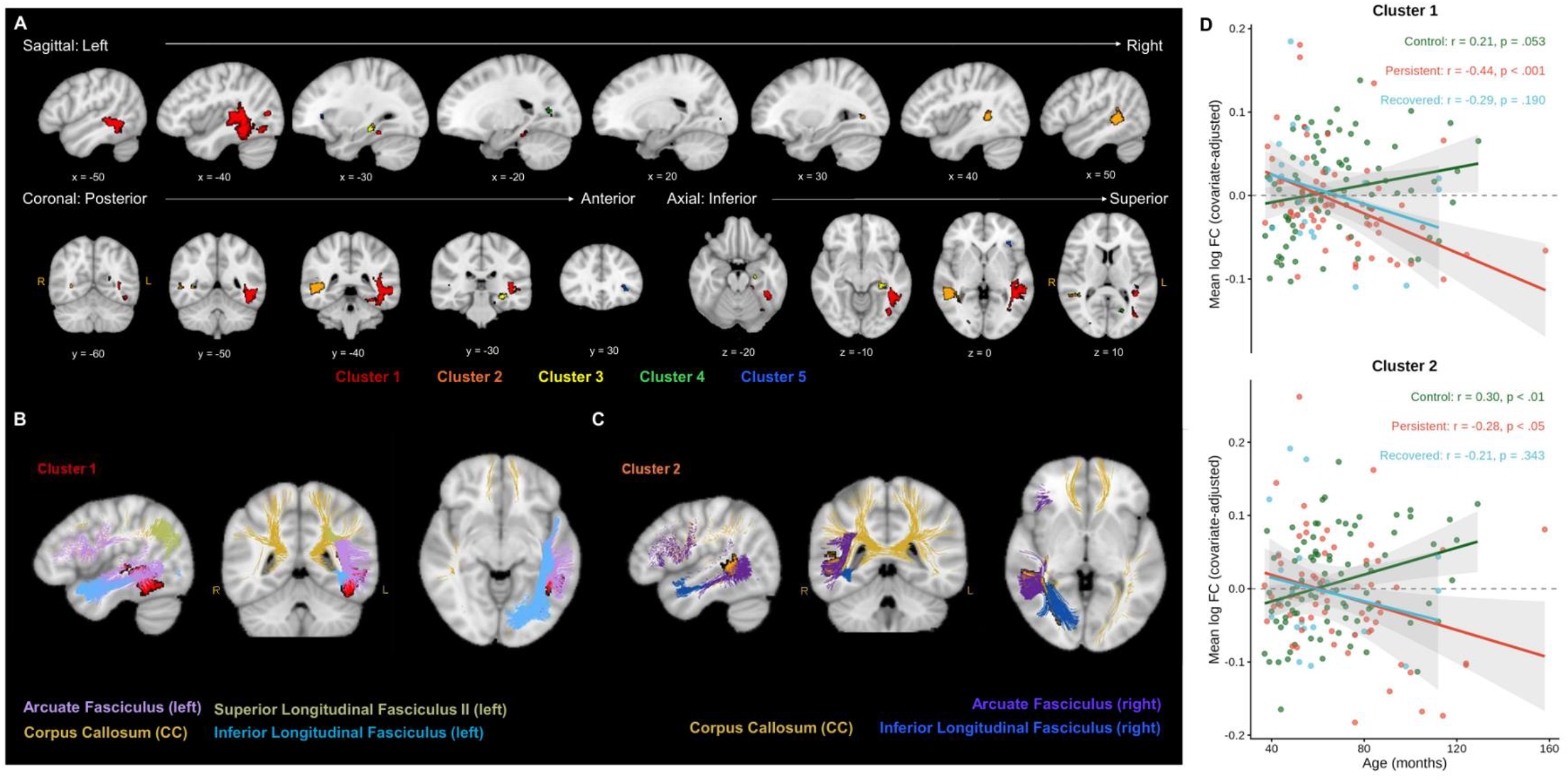
Children who stutter show reduced age-related increases in fiber cross-section across bilateral white matter pathways. A. Whole-brain Group x Age effects in FC (CWS < controls; *p*_unc_ < .001, cluster extent > 10 voxels). B – C. Fiber populations within the two largest clusters, demonstrating overlap with multiple white matter pathways. Cluster 1 (Peak MNI coordinates: −52, −63, −19 (mm)) was predominantly associated with the left arcuate fasciculus (AF), and Cluster 2 (Peak MNI coordinates: 15, - 61, −14 (mm)) with the right AF. D. Age-related changes of covariate-adjusted log FC for Clusters 1 and 2, illustrating greater age-related increases in controls than in CWS. Detailed anatomical overlap is reported in Supplemental Table 2.

Within-group associations between age and log FC were examined descriptively within these clusters. In typically developing children, significantly positive correlations between age and FC were observed for cluster 2 (r = 0.30, *p* <.005) and cluster 5 (r = 0.33, *p* <.005). For the persistent stuttering subtype, significantly negative correlations were observed for cluster 1(r = −0.44, *p* <.001), cluster 3 (r = −0.42, *p* <.001), and cluster 5 (r = −0.46, *p* <.001). No significant associations were observed in the recovered subgroup. Stuttering severity scores did not show significant correlations with Fiber cross-section across either persistent or recovered stuttering subgroups.

### 3.2 Tract of Interest (TOI) Analyses

For significant whole-brain results in group and group x age interactions, we applied an a priori tract-based analysis to investigate how specific tracts of interest are implicated and in what way. Supplemental Figure 3 and Figure 3 list the tracts of interest with significant Fixels identified at *p_FWE_* <.05 for Group effects (CWS < Controls) and group x age Interaction (CWS < Controls), respectively.

**Figure 3.**
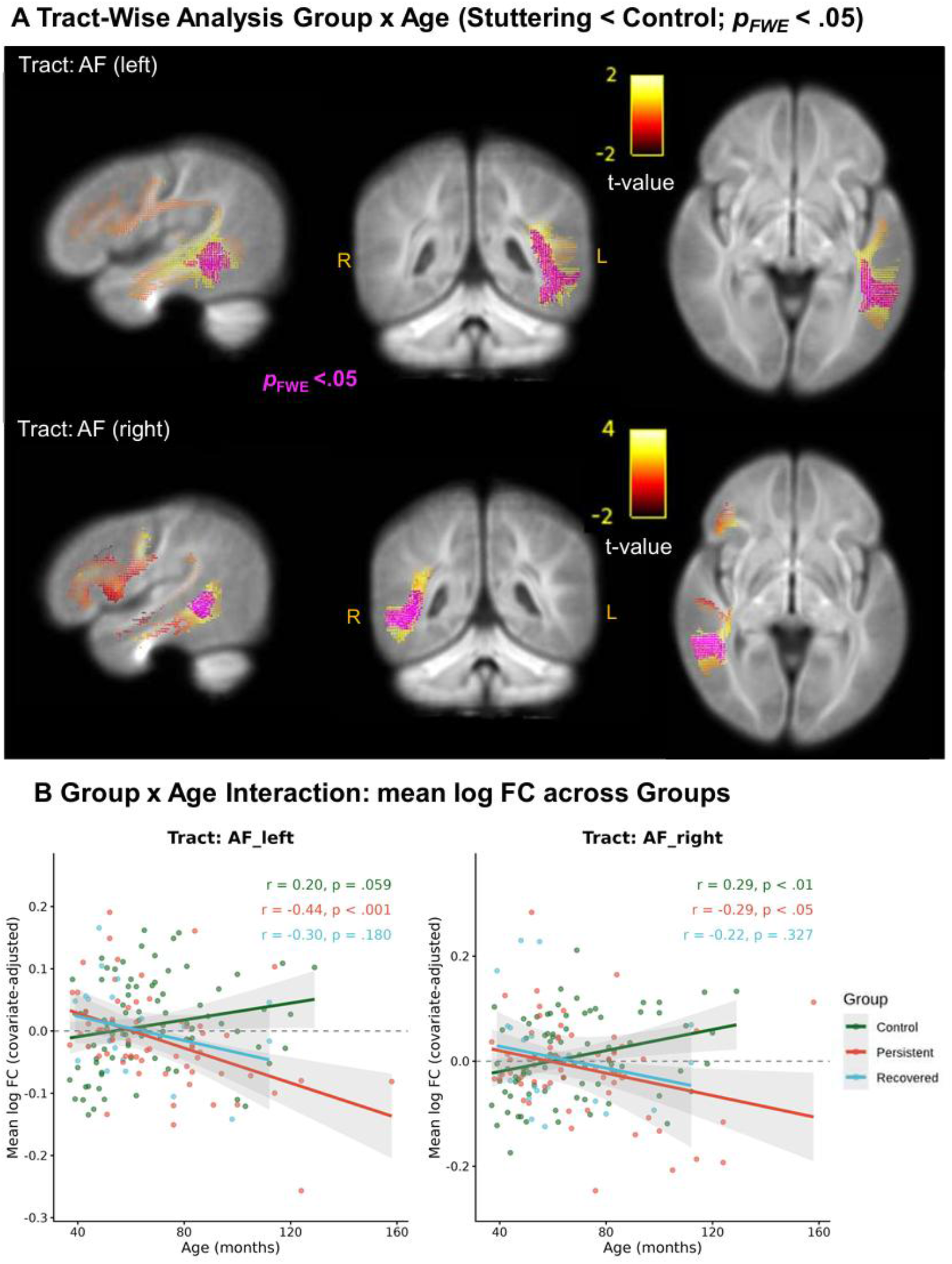
Bilateral arcuate fasciculus shows divergent age-related fiber cross-section trajectories in children who stutter. A. Tract-of-interest analysis showing Fixels with a Group × Age interaction (CWS < controls) at the within-tract FWE-corrected threshold (*p***_FWE_** < .05; magenta), overlaid on the corresponding statistical maps. Effects were observed in the left and right arcuate fasciculus (AF) and right corticospinal tract (CST). Following BH-FDR correction across the 11 tract-of-interest analyses, effects remained significant in the bilateral AF (*q* = .0187). B. Age-related trajectories of covariate-adjusted log FC within significant Fixels, illustrating greater age-related increases in controls relative to CWS.

#### 3.2.1 Group effects: significant differences in tracts, and relationship with stuttering severity

Tract-wise GLMs identified group differences in FC reaching within-tract FWE-corrected significance (*p*FWE < .05) in several a priori tracts (Supplemental Figure 3). No a priori tract showed significantly greater FC in CWS relative to controls (CWS > controls) at the within-tract FWE-corrected threshold. Lower FC in CWS relative to controls (CWS < controls) was observed in the left arcuate fasciculus (AF), left anterior thalamic radiation (ATR), left superior thalamic radiation (STR), left superior longitudinal fasciculus III (SLF III), and bilateral corticospinal tracts (CST). However, these effects did not remain significant following BH-FDR correction across the 11 tract-based analyses (*q* > .05). Stuttering severity scores were not significantly correlated with FC within either the persistent or recovered stuttering subgroups. Full tract-wise results, including minimum FWE-corrected *p* values, the extent of FWE-significant Fixels, and BH-FDR-adjusted *q* values, are reported in Supplemental Tables 3 and 4.

Anatomical localization of the Fixel-wise effects using the JHU and ICMB white matter atlases showed that tracts showing significantly lower FC for CWS relative to controls were localized to the left corticospinal tract, the left cerebral peduncle, and to the posterior limb of the left internal capsule. Within-group descriptive associations between age and log FC showed significantly positive correlations between age and FC in the typical control group for Left CST (r = 0.28, *p* <.05), right CST (r = 0.31, *p* <.005), and left STR (r = 0.27, *p* <.01). For the persistent stuttering subtype, significantly positive correlations were observed for left CST (r = 0.52, *p* <.05). No significant associations were observed in the persistent stuttering subgroup. Stuttering severity scores did not show significant correlations with Fiber cross-section across either persistent or recovered stuttering subgroups.

#### 3.2.2 Differences in age-related associations between groups: significant differences in tracts, within-group associations, and relationship with stuttering severity

Tract-wise GLMs revealed significant Group × Age interactions in FC at the within-tract FWE-corrected threshold (*p*FWE < .05) across several a priori tracts (Figure 3 and Supplemental Figure 6; Table 3 and Supplemental table 5). Group × Age interactions, reflecting a greater age-related increase in FC in controls relative to CWS (CWS < controls), were observed bilaterally in the arcuate fasciculus (AF) and in the right corticospinal tract (CST). Following additional BH-FDR correction across the 11 tract-based analyses, the interactions in the left and right AF remained significant (both *q* = .0198), whereas the right CST interaction did not (*q* = .1115). Atlas-based anatomical correspondence using the JHU and ICBM White Matter Atlas indicated partial overlap with these and other tracts (e.g., inferior fronto-occipital fasciculus, forceps major, posterior thalamic radiation, sagittal stratum, retrolenticular internal capsule), particularly in regions of known fiber crossing, limiting tract-specific interpretation. Within-group descriptive analysis showed that this interaction was driven by the persistent stuttering subgroup for left AF, which exhibited a significant negative association between mean log fiber cross-section (FC) and age (r = −0.44, *p* <.001). For the right AF, the typical control group exhibited a significant positive association (r = 0.29, *p* < .05), and the persistent stuttering subgroup exhibited a significant negative association (r = −0.29, *p* < .05) between mean log fiber cross-section (FC) and age.

**Table 3.**
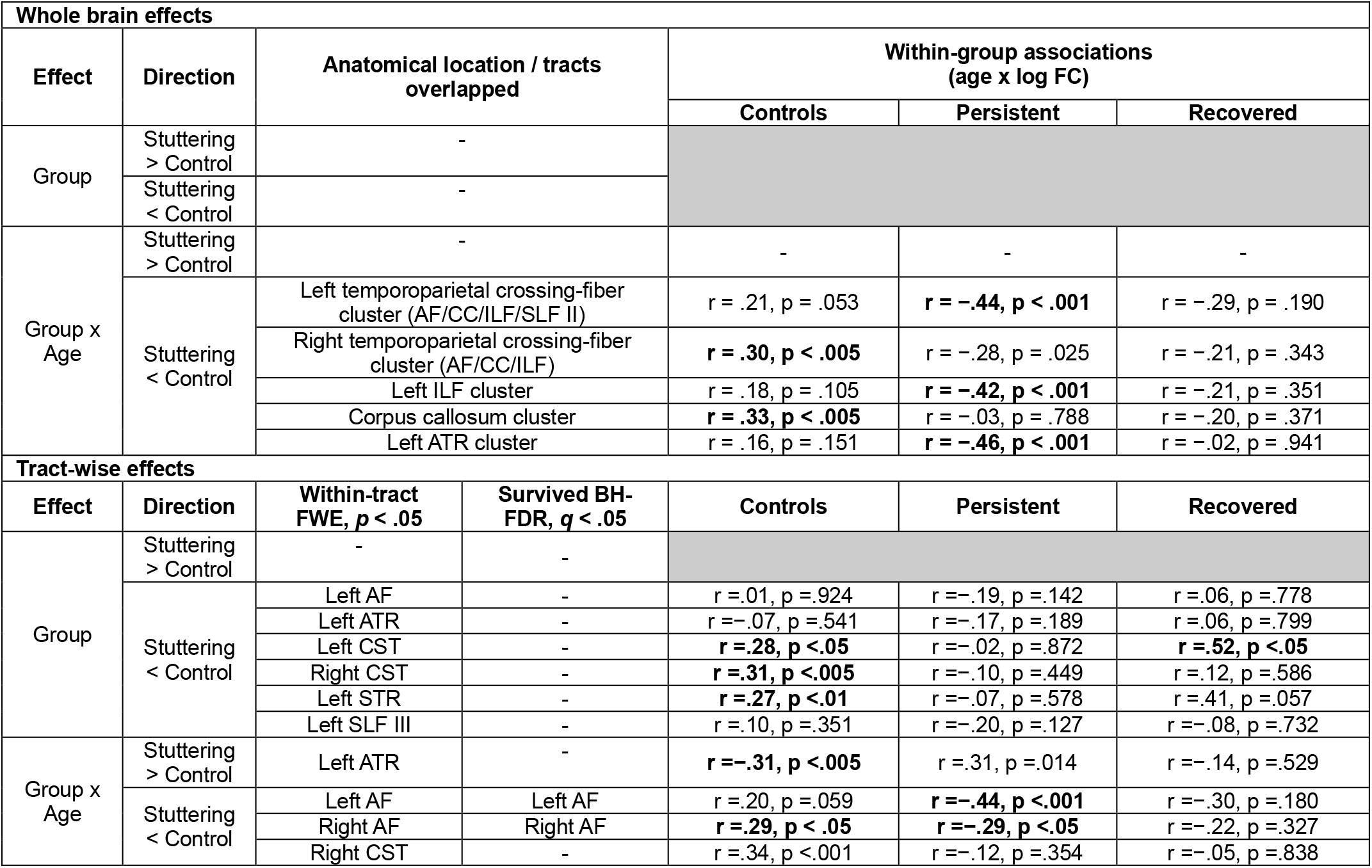
Summary of whole-brain and tract-wise fiber cross-section findings.

**Table 4.** Tract-of-Interest (TOI) Fixel-wise analysis of Group × Age interaction (CWS < CWNS).

| Rank | Tract | Min. FWE-corrected *p* | FWE-significant Fixels, n (%) | *q*<sub>BH_FDR</sub> |
| --- | --- | --- | --- | --- |
| 1 | Arcuate Fasciculus (L) | **.0020\*** | **2037 (9.08%)** | **.0198\*** |
| 2 | Arcuate Fasciculus (R) | **.0036\*** | **1033 (4.60%)** | **.0198\*** |
| 3 | Corticospinal Tract (R) | **.0304\*** | **97 (2.52%)** | .1115 |
| 4 | Superior Thalamic Radiation (R) | .0564 | 0 (0%) | .1214 |
| 5 | Superior Longitudinal Fasciculus III (R) | .0632 | 0 (0%) | .1214 |
| 6 | Corpus Collosum | .0662 | 0 (0%) | .1214 |
| 7 | Superior Thalamic Radiation (L) | .1112 | 0 (0%) | .1560 |
| 8 | Anterior Thalamic Radiation (L) | .1272 | 0 (0%) | .1560 |
| 9 | Superior Longitudinal Fasciculus III (L) | .1276 | 0 (0%) | .1560 |
| 10 | Corticospinal Tract (L) | .1742 | 0 (0%) | .1916 |
| 11 | Anterior Thalamic Radiation (R) | .2644 | 0 (0%) | .2644 |
| **Note.** *p* values are fixel-wise FWE-corrected within each tract. The minimum FWE-corrected *p* value from each of the 11 primary tract analyses was subjected to Benjamini–Hochberg FDR correction across tracts. *q*<sub>BH-FDR</sub> denotes the BH-FDR-adjusted tract-level *p* value. Bold values indicate *p* < .05 for within-tract FWE results and *q* < .05 for across-tract FDR-corrected results. |  |  |  |  |
| **BH-FDR alpha:** .05; **Largest rejected BH rank:** 2; **BH raw-*p* rejection cutoff:** .00360; **Equivalent 1-*p* cutoff:** .9964. |  |  |  |  |

Group × Age interaction, reflecting a greater age-related increase in FC in CWS relative to controls (CWS > controls), was observed in the left anterior thalamic radiation (ATR) at the within-tract FWE-corrected threshold; however, this effect did not survive BH-FDR correction across tracts (*q* = .2178; Supplemental Figure 5). Thus, the bilateral AF effects represented the Group × Age interactions that remained significant following both within-tract FWE and across-tract multiple-comparison correction.

Atlas-based anatomical correspondence using the JHU White Matter Tractography Atlas indicated that the significant cluster (CWS > controls) overlapped primarily with the left anterior thalamic radiation. Additional correspondence with the JHU ICBM-DTI-81 White Matter Labels Atlas suggested minor overlap with the left anterior limb of the internal capsule and negligible overlap with the left posterior limb of the internal capsule. Within-group descriptive analysis showed that this interaction was primarily driven by the typical control group for left ATR, which exhibited a significant negative association between mean log fiber cross-section (FC) and age (r = −0.31, *p* <.005).

### 3.3 FA versus FBA results comparison

To examine the spatial correspondence between the current Fixel-based findings and those obtained using a conventional diffusion tensor metric, we performed a complementary fractional anisotropy (FA) analysis using Tract-Based Spatial Statistics (TBSS) in the same dataset. No TBSS-based FA analysis survived family-wise error correction at *p* < .05. At an uncorrected threshold of *p* < .001, the FA findings for the group contrast (CWS < controls) overlapped spatially with the FC findings in the left corticospinal tract (CST). For the Group × Age interaction (CWS < controls, FC and FA findings showed partial spatial overlap in posterior regions of the left and right arcuate fasciculus (AF). No clear spatial correspondence between FC and FA findings was observed for the CWS > controls group or group × Age contrasts. These comparisons were exploratory and were performed to characterize the degree of spatial correspondence between the two diffusion-analysis approaches rather than as a formal test of agreement between FA and FC.

## 4. Discussion

### 4.1 Group differences in Fiber Cross Section in Children Who Stutter compared to typically developing children

In this study, we examined differences in microstructural (fiber density, FD) and macrostructural (fiber cross-section, FC) properties of white matter, as well as their combined index (FDC), between CWS and typically developing children. Among these measures, only FC showed statistically significant group effects and group-by-age interaction in the whole-brain analysis, while FD and FDC did not significantly differ between groups.

Whole-brain group differences in FC were localized to major projection pathways connecting cortical and subcortical regions, with additional involvement of association fibers supporting intrahemispheric connectivity. Specifically, increased FC in CWS relative to typically developing children was anatomically localized to regions within right anterior thalamic radiation (ATR), right inferior fronto-occipital fasciculus (IFOF), and right anterior corona radiata. The tract-of-interest analyses did not identify corresponding increases in FC in CWS at the within-tract FWE-corrected threshold. Thus, these increases appear to represent spatially localized whole-brain effects rather than robust tract-wide differences. The whole-brain spatial pattern may reflect altered connectivity across more than one functional system: the ATR supports thalamocortical relay circuits implicated in processing speed and executive function (Ferris et al., 2022), while the IFOF is increasingly recognized for a role in executive control and attentional processing (Giampiccolo et al., 2025). These findings may therefore relate not only to motor timing and initiation, as previously discussed, but also to executive and attentional control processes relevant to speech planning.

Increased FC is generally understood to reflect larger cross-sectional areas of a fiber bundle, indicating macrostructural expansion of the bundle or altered extracellular architecture within the pathway (D. A. Raffelt et al., 2017). However, the specific biological substrate cannot be determined by FC alone. Given these considerations, the right lateralized pattern of increased FC observed in the current study may reflect a form of structural reorganization, although the current data cannot adjudicate between competing accounts of the root cause; it may be due to either compensatory reorganization, maladaptive integration, or delayed pruning of the fiber bundles. This right lateralization pattern has previously been reported in adults who stutter, but not in children who stutter, in previous studies exploring traditional diffusion and structural connectivity measures (Foundas et al., 2003). White matter asymmetries have also been linked more broadly to the lateralization of other functions, such as language production and spatial attention (Zhu et al., 2025). The presence of this pattern in children provides preliminary evidence that such reorganization, if compensatory, may be present at a younger age than initially anticipated. However, the current design cannot distinguish between competing mechanistic accounts, and thus, these interpretations should be considered speculative. The structural expansion in fiber bundles may not necessarily indicate efficient compensatory processing and may instead indicate an altered suboptimal integration that is maladaptive. It could also be that proper pruning (Peter R., 1979; Tau & Peterson, 2010) of these right-hemispheric fibers is delayed in the development trajectory of children who stutter (Beal et al., 2013, 2015). Further work exploring FC trajectories longitudinally may provide additional details to resolve exact mechanisms at play.

On the other hand, decreased FC is generally understood to reflect a smaller cross-sectional area of a fiber bundle, potentially indicating fewer axons, reduced axonal packing, or a narrower overall pathway caliber (D. A. Raffelt et al., 2017). In the current study, at a whole-brain level, reductions in FC in CWS were anatomically localized to the left corticospinal tract, with extensions into the left cerebral peduncle and posterior limb of left internal capsule, indicating alterations in projection pathways, alongside more focal involvement of descending motor pathways responsible for motor execution. Tract-of-interest analyses additionally identified lower FC in CWS within the left anterior thalamic radiation, left superior thalamic radiation, and bilateral corticospinal tracts at the within-tract FWE-corrected threshold. However, none of these tract-level group effects remained significant following additional BH-FDR correction across the 11 a priori tract analyses. Together with the whole-brain findings, the spatial pattern is consistent with altered macrostructural organization of motor and thalamocortical pathways in CWS, although the present data should be interpreted more cautiously as the effects did not survive correction across tracts. Collectively, these findings may be consistent with a model of disrupted left-hemisphere motor pathways, potentially accompanied by compensatory but potentially inefficient recruitment of right-hemisphere networks. However, the current data cannot confirm this account over alternative explanations, and functional interpretations (e.g., links to auditory-motor integration) remain speculative.

### 4.2 Changes in group differences by age in Fiber Cross Section in children who stutter compared to typically developing children

The most robust tract-wise developmental differences were observed bilaterally in the arcuate fasciculus (AF). Group × Age interactions in both the left and right AF survived within-tract FWE correction as well as additional BH-FDR correction across the 11 a priori tract analyses. In both tracts, FC increased with age to a greater extent in typically developing controls than in CWS. Descriptive within-group analyses further indicated positive age-related FC trajectories in controls, whereas children with persistent stuttering showed negative associations between age and FC. Together, these findings suggest divergence in the developmental trajectory of bilateral arcuate fiber-bundle cross-section in childhood stuttering, particularly among children with persistent symptoms.

The bilateral AF forms a major component of the dorsal language network and supports communication between temporal and frontal cortical regions involved in speech and language processing (Hickok et al., 2011; Brauer et al., 2013). Its involvement is therefore consistent with atypical development of structural pathways supporting auditory-motor integration. Importantly, the bilateral nature of the present findings suggests that developmental differences are not restricted to left hemisphere speech pathways. Rather, the trajectory of fiber-bundle macrostructure appears to diverge between CWS and controls in homologous association pathways across both hemispheres. Longitudinal data will be necessary to establish whether these cross-sectional age effects reflect altered rates of fiber-bundle growth, differences in developmental timing, or other forms of structural reorganization.

Additional Group × Age effects reached the within-tract FWE-corrected threshold but did not survive correction across the 11 tract analyses. A negative interaction was observed in the right corticospinal tract (CST), suggesting reduced age-related FC increases in CWS relative to controls, whereas a positive interaction was observed in the left anterior thalamic radiation (ATR), indicating a greater age-related increase in FC in CWS (Supplemental Figure 5). The latter effect was primarily associated with the persistent stuttering subgroup, indicating atypical maturation of thalamocortical pathways in this group. Although these findings may point to additional developmental differences involving motor execution and thalamocortical pathways, their failure to survive across-tract BH-FDR correction warrants cautious interpretation and independent replication.

The left ATR finding is nevertheless of potential interest given the role of thalamocortical pathways in cognitive and motor functions. The ATR connects the anterior and mediodorsal thalamic nuclei with medial and dorsolateral prefrontal cortex, and structural and functional disruption along this pathway has been linked to deficits spanning attention, working memory, executive control (Schumacher et al., 2019), and language processing (Stockert et al., 2025). This broader functional profile is consistent with a substantial behavioral literature reporting weaker inhibitory control, attentional regulation, and executive function in children who stutter relative to typically speaking peers (J. D. Anderson & Ofoe, 2019; Ofoe et al., 2018). An atypical developmental trajectory in this pathway could therefore have relevance beyond speech motor timing and initiation. However, whether the observed FC increase reflects compensatory structural reorganization, altered developmental scaling, or another mechanism cannot be determined from the present cross-sectional data.

Prior longitudinal work reports that persistent and recovered CWS show diverging white matter growth trajectories relative to controls, including in tracts overlapping the anterior thalamic radiation, with some tracts showing atypical increases alongside others showing reductions (Chow & Chang, 2017a). Separately, connectivity between subcortical and frontal regions (e.g., cerebellar-orbitofrontal circuits) has been shown to scale inversely with stuttering severity, consistent with a compensatory role for specific cortico-subcortical pathways in more fluent speakers who stutter (Sitek et al., 2016). Whether the FC increase we observe specifically reflects greater recruitment of effortful, top-down control during speech production, as opposed to a more general compensatory mechanism, is not established by the current data and remains speculative. While atlas-based correspondence localized the effect predominantly to the anterior thalamic radiation, with minor extension into the anterior limb of the internal capsule, the spatial specificity of Fixel-based measures suggests that these findings reflect focal alterations within thalamocortical circuitry rather than widespread voxel-wise changes. Together, these results point to atypical development of thalamocortical networks as a key feature of persistent stuttering, potentially contributing not only to disruptions in the timing and initiation of speech, but also to broader difficulties in cognitive control and attentional regulation that have been reported behaviorally in this population.

Overall, the Group × Age findings provide strongest evidence for divergent development of bilateral arcuate fiber-bundle macrostructure in CWS relative to typically developing controls. The additional within-tract findings suggest that developmental differences may extend to corticospinal and thalamocortical pathways, but these effects were not robust to correction across the full set of a priori tract analyses. This distinction narrows the principal tract-wise developmental finding to bilateral AF while identifying the other pathways as secondary findings warranting further investigation.

### 4.3 Complementary insights from Fixel-based analysis and fractional anisotropy

Fixel-based analysis provides three complementary information to conventional diffusion tensor metrics such as fractional anisotropy (FA) by characterizing individual fiber populations within a voxel and distinguishing microstructural and macrostructural properties of white matter. Fiber Density (FD) reflects the intra-axonal volume fraction within a given fiber population. Fiber cross-section (FC), by contrast, reflects the overall cross-sectional area of a fiber bundle relative to the study population, capturing bundle-level expansion or atrophy rather than the density of axons packed within it. The combined metric, fiber density and cross-section (FDC), integrates both properties and reflects the total capacity of a bundle to convey information, though at the cost of specificity as to which property is driving an observed effect (D. A. Raffelt et al., 2017).

Moreover, because FBA is built on constrained spherical deconvolution rather than the single-tensor model underlying conventional DTI, it resolves multiple crossing fiber populations within a voxel, avoiding the interpretive ambiguity inherent to single-tensor metrics such as FA - a limitation affecting a substantial proportion of white matter voxels (Jeurissen et al., 2013; D. A. Raffelt et al., 2017). This approach therefore offers a unique opportunity to quantify the contribution of individual white matter tracts to group differences in white matter integrity measures.

The complementary FA analysis in the current study demonstrated partial spatial correspondence with the FC findings. For the CWS > controls Group effect, FC and FA findings overlapped within the left corticospinal tract, while the CWS> controls Group × Age effects showed overlap in posterior regions of the bilateral arcuate fasciculus. This convergence is consistent with previous DTI studies reporting white-matter differences involving speech and motor pathways in developmental stuttering (Chang et al., 2008; Chang & Zhu, 2013; Chang et al., 2015; Chow & Chang, 2017a). However, the correspondence was incomplete, and the two measures should not be expected to produce equivalent spatial patterns because they quantify different properties of white-matter organization.

The FC findings therefore provide information that cannot be inferred directly from FA. In particular, the bilateral arcuate Group × Age effects identified using FC indicate developmental differences in the macrostructural cross-section of specific fiber populations. The ability of FBA to model multiple fiber populations within crossing-fiber regions provides greater anatomical specificity for attributing these effects to particular fiber pathways. Nevertheless, the present comparison does not establish that previously reported FA effects were caused by, or independent of, crossing-fiber contributions, nor does spatial correspondence between FA and FC establish a common underlying biological mechanism.

Prior DTI studies have interpreted reduced FA together with changes in radial and axial diffusivity as potentially reflecting differences in white-matter microstructure, including myelination (Cykowski et al., 2010). FC addresses a different aspect of white-matter organization and therefore neither confirms nor contradicts such interpretations. Instead, the present results suggest that developmental stuttering may also involve differences in fiber-bundle macrostructure. Future longitudinal and multimodal studies combining fiber-specific diffusion measures with more direct measures of tissue microstructure will be necessary to determine how macrostructural bundle development, axonal organization, and myelination jointly contribute to the white-matter differences observed in developmental stuttering.

### 4.4 FC changes in the absence of FD and FDC differences suggest tract-level macrostructural alterations

A notable finding in the present study is the dissociation between Fixel-based metrics, with significant differences observed in FC but not in FD or FDC. This pattern contrasts with other neurodevelopmental findings, where microstructural differences (e.g., FD) are more commonly reported, often in the absence of or with FC-based macrostructural changes. In several neurodevelopmental disorders, such as autism spectrum disorder (Dimond et al., 2019; Kirkovski et al., 2024), attention deficit/hyperactivity disorder (Hyde et al., 2021; Thomson et al., 2022), and cerebral palsy (Crotti et al., 2025; Tsai et al., 2026), alterations in FD have been observed with relatively stable or sometimes significant effects in FC across groups, suggesting microstructural disruption as a primary feature. FC changes remain more heterogeneous across disordered cohorts. Many neurodevelopment cohorts show positive associations between age and FD. But more specific increases in FC across specific pathways (Dimond et al., 2020; Genc et al., 2018). The current results indicate that CWS mainly exhibit differences at a macrostructural level, reflected in altered FC, while microstructural properties indexed by FD remain largely unchanged. This may suggest that white matter differences in children who stutter are driven by large-scale tract morphology rather than by fiber-specific density changes.

One possible interpretation is that these FC differences reflect neurodevelopmental adaptation rather than primary deficits. Given that our sample spans both preschool (3 – 6 years) and school-age (6 – 13 years) children, the observed pattern may reflect some combination of early developmental differences and subsequent compensatory processes, though our linear age model cannot dissociate these possibilities. For example, increased FC could reflect processes such as delayed pruning or altered tract specialization, leading to broader or less refined white matter pathways. From a functional perspective, larger or less specialized pathways may reflect increased processing demands or reduced efficiency within relevant networks. This is consistent with evidence that atypical or delayed white matter refinement is associated with less efficient information processing (Scantlebury et al., 2014; Yeatman et al., 2012) and with altered neural signaling in other neurodevelopmental populations (Hanson et al., 2025). However, whether these changes represent a primary developmental difference or a compensatory response to early deficits remains an open question that warrants further investigation. It should also be noted that the current study modeled age linearly, such that nonlinear effects were not tested. Thus, a clear bimodal distribution between the two age groups was not identified.

### 4.5 Limitations

The current study contains several limitations with regard to the sample size as well as diffusion acquisition parameters that could affect the results adversely. Similar to other clinical cohorts studied, our group sizes were relatively small compared to large-scale open datasets. This could affect statistical significance and power to note small-to-moderate effects. The standard b-values widely used in FBA studies range from 1000 – 3000 s/mm^2^ (single-shell or multi-shell), and multi-shell is preferred (i.e., multiple b-value acquisitions are present, e.g., b = 1000 + 2000 + 3000). It is possible to complete FBA with lower b-values that are slightly lower and with single-shell acquisitions, but the Fiber orientation density (FOD) angular resolution decreases, effects of partial volumes increase, and the combined effect of these causes the Fiber Density estimates to become less reliable (Genc et al., 2020). Thus, diffusion acquisition parameters in the current study may have limited sensitivity for accurately estimating fiber density (FD), as reliable FOD reconstruction typically benefits from higher b-values (≥2000 s/mm^2^). As a result, null findings in FD should be interpreted cautiously.

### 4.6 Future Directions

Future directions should extend this work to adult populations who stutter to determine whether the observed macrostructural patterns persist through development or reflect developmental reorganization. Longitudinal and interventional designs will be particularly important for characterizing how white matter properties assessed via Fixel-based metrics evolve over time (Genc et al., 2018; Kelly et al., 2020) In this context, FBA offers important advantages over conventional diffusion tensor imaging metrics, as it enables the separation of fiber-specific properties within crossing fiber regions and provides a clear characterization between macro- vs. micro-structure of white matter organization. These properties make FBA particularly well-suited for applications involving predictive biomarkers for neuromodulation as well as neuromodulation target selection. Although the current study was cross-sectional and based on a modest sample size, observed effects were consistent with prior literature, supporting the robustness of the findings. Future studies should therefore consider leveraging FBA in longitudinal and intervention-based frameworks to better characterize developmental trajectories and treatment-related changes, particularly in the context of neuromodulatory approaches.

## 5. Conclusion

This study provides the first application of Fixel-based analysis (FBA) to developmental stuttering and demonstrates that white-matter differences in children who stutter are primarily detectable in fiber cross-section (FC), a measure of fiber-bundle macrostructure, rather than fiber density (FD). Whole-brain and tract-of-interest analyses identified differences in FC across association, projection, and thalamocortical pathways. Most notably, developmental differences in the bilateral arcuate fasciculus remained significant following both within-tract FWE correction and additional correction for multiple comparisons across tracts, providing the strongest evidence for divergent development of fiber-bundle macrostructure in pathways supporting auditory-motor integration. Other thalamocortical and corticospinal effects provide additional, but less robust, evidence of altered white-matter development in childhood stuttering.

FBA provided complementary information to conventional tensor-based diffusion measures by enabling fiber-specific characterization of white matter, particularly in regions containing crossing fiber populations. The partial spatial correspondence between FC and FA findings suggests that the two approaches capture related but non-equivalent features of white-matter organization. Together, these findings highlight altered development of fiber-bundle macrostructure as a feature of developmental stuttering and demonstrate the value of fiber-specific diffusion approaches for characterizing its neurodevelopmental basis. Longitudinal and independent replication studies will be important for determining how these structural trajectories relate to persistence and recovery.

## 6. Code and Data Sharing Statement

Analysis code used in this study, including scripts for Fixel-based analysis, statistical modeling, tract-of-interest analyses, and multiple-comparison correction, is publicly available on GitHub at [https://github.com/HasiniWeerathunge/FixelBasedAnalysis]. The repository also includes documentation describing the analysis workflow and software dependencies. De-identified data supporting the findings of this study may be made available upon reasonable request and subject to institutional and ethical requirements governing participant data sharing.

## Author contributions

H.W. and S.E.C. conceived and designed the study. F.H., H.M.C., A.J., and M.A. guided the study design and troubleshooting of the analyses. H.W. performed the image processing, statistical analyses, and visualization. H.W., Y.L., and S.E.C. interpreted the findings. H.W. drafted the initial manuscript. S.E.C. and Y.L. supervised the project. S.E.C. provided project resources and acquired funding. All authors critically revised the manuscript for intellectual content and approved the final version.

## Acknowledgements

We thank Megan Sheppard, speech-language pathologist in the Speech Neurophysiology Lab at the University of Michigan, for her valuable contributions to participant recruitment, data collection, speech and language assessments, data curation, and assistance with measurement-related and data queries. We also thank the research assistants and staff of the Speech Neurophysiology Lab for their contributions to participant recruitment, data collection, and study coordination. Finally, we extend our sincere gratitude to all children who participated in this study and to their parents and families for their time, commitment, and support.

## Funding

This work was supported by the National Institute on Deafness and Other Communication Disorders (NIDCD) of the National Institutes of Health (NIH; R01DC011277 to S.-E.C.).

## Notes

### Competing Interest Statement

The authors have declared no competing interest.

