## Supplemental Tables and Figures for "Fixel-wise differences in White Matter Fiber-bundle Cross-section identified in Children who Stutter"

**Supplemental Table 1. Participant sex distribution and age characteristics stratified by diffusion MRI acquisition cohort.** Age is reported in months as mean (SD; standard deviation) and rage separately for control and stuttering groups within cohorts 1 and 2. Cohort 1 and 2 correspond to the two diffusion acquisition protocols used in the study.

| Acquisition cohort | Group | N | Sex | Age, months Mean (SD) | Range |
|---|---|---|---|---|---|
| **Cohort 1** | Control | 40 | 21 male / 19 female | 77.18 (25.32) | 39 – 129 |
| | Stuttering | 42 | 27 male / 15 female | 74.69 (27.20) | 37 – 158 |
| **Cohort 2** | Control | 47 | 23 male / 24 female | 58.21 (12.78) | 37 – 83 |
| | Stuttering | 42 | 24 male / 18 female | 61.14 (17.55) | 38 – 124 |

**Supplemental Table 2. Anatomical localization and white-matter tract overlap of clusters showing CWS < controls Group x Age interaction in log-transformed fiber cross-section (logFC).**

| Cluster | Extent (voxels) | Peak MNI (x,y,z)(mm) | JHU ICBM-DTI-81 labels (% cluster) | JHU probabilistic tracts ≥25% (% cluster) | Study-specific tract overlap (% cluster fixels) |
|---|---|---|---|---|---|
| **1** | 3025 | (−52, −63, −19) | L posterior thalamic radiation 4.69%;<br>L sagittal stratum 3.67%;<br>L SLF 3.60%;<br>L retrolenticular internal capsule 1.39% | L ILF 17.79%;<br>L IFOF 7.64%;<br>L SLF 5.69%;<br>L temporal SLF 5.19% | L AF 31.49%;<br>L ILF 7.79%;<br>CC 6.72%;<br>L SLF-II 0.03% |
| **2** | 1072 | (15, −61, −14) | R posterior thalamic radiation 14.83%; R SLF 6.34% | R IFOF 10.26%;<br>forceps major 3.36%;<br>R temporal SLF 1.21%;<br>R SLF 0.65%;<br>R ILF 0.19% | R AF 42.45%;<br>CC 5.91%;<br>R ILF 3.60% |
| **3** | 161 | (−48, −57, −20) | L fornix (cres)/stria terminalis 3.11% | None ≥25% | L ILF 0.26% |
| **4** | 47 | (−36, −80, −6) | Splenium of corpus callosum 10.64% | None ≥25% | CC 3.66% |
| **5** | 44 | (−49, −12, −13) | None | None ≥25% | L ATR 8.40% |

**Note:** Clusters identified at $p_{unc} < .001$ with a minimum extent of 10 voxels, with overlapping anatomical atlases or study-specific tracts are presented. Cluster extent and center-of-mass MNI coordinates are reported. Anatomical localization used the JHU ICBM-DTI-81 labels and JHU probabilistic tractography atlases (≥25% probability). Overlap percentages represent the proportion of cluster voxels within atlas regions or cluster Fixels intersecting study-specific tracts. Two additional clusters that fit the 10 voxel criteria (6 and 7) were omitted due to absent overlapping. Cluster characterization was descriptive; whole-brain inference was based on connectivity-based Fixel enhancement with family-wise error correction. **SLF:** Superior Longitudinal Fasciculus; **ILF:** Inferior Longitudinal Fasciculus; **IFOF**: Inferior Frontal-Occipital Fasciculus; **AF**: Arcuate Fasciculus; **CC**: Corpus Callosum; **ATR**: Anterior Thalamic Radiation; **L**: Left; **R**: Right.

**Supplemental Table 3. Tract-of-Interest (TOI) Fixel-wise analysis of the Group main effect (CWS > controls).** Results from connectivity-based Fixel enhancement (CFE) analyses performed separately within 11 a priori white-matter tract masks. $p_{FWE}$ represents the minimum Fixel-wise family-wise error-corrected p-value within each tract, and *% FWE-significant Fixels* represents the percentage of Fixels within the tract with FWE-corrected p < .05. BH-FDR correction was subsequently applied across the 11 tract-level minimum FWE p-values, with q < .05 considered significant.

| **Rank** | **Tract** | **Min. FWE-corrected *p*** | **FWE-significant Fixels, n (%)** | $q_{BH\_FDR}$ |
|---|---|---|---|---|
| 1 | Anterior Thalamic Radiation (R) | .0798 | 0 (0%) | .4560 |
| 2 | Superior Longitudinal Fasciculus III (R) | .1526 | 0 (0%) | .4560 |
| 3 | Arcuate Fasciculus (R) | .1652 | 0 (0%) | .4560 |
| 4 | Corpus Callosum | .1658 | 0 (0%) | .4560 |
| 5 | Superior Thalamic Radiation (R) | .6014 | 0 (0%) | .9682 |
| 6 | Superior Thalamic Radiation (L) | .7050 | 0 (0%) | .9682 |
| 7 | Superior Longitudinal Fasciculus III (L) | .7956 | 0 (0%) | .9682 |
| 8 | Anterior Thalamic Radiation (L) | .8244 | 0 (0%) | .9682 |
| 9 | Arcuate Fasciculus (L) | .8968 | 0 (0%) | .9682 |
| 10 | Corticospinal Tract (L) | .9520 | 0 (0%) | .9682 |
| 11 | Corticospinal Tract (R) | .9682 | 0 (0%) | .9682 |
| **Note.** p values are fixel-wise FWE-corrected within each tract. The minimum FWE-corrected p value from each of the 11 primary tract analyses was subjected to Benjamini–Hochberg FDR correction across tracts. qBH-FDR denotes the BH-FDR-adjusted tract-level p value. Bold values indicate p < .05 for within-tract FWE results and q < .05 for across-tract FDR-corrected results. | | | | |
| No tract reaches within-tract FWE significance or survives BH-FDR. | | | | |

**Supplemental Table 4. Tract-of-Interest (TOI) fixel-wise analysis of the Group main effect (CWS < controls).**

| Rank | Tract | Min. FWE-corrected *p* | FWE-significant Fixels, n (%) | $q_{BH\_FDR}$ |
|---|---|---|---|---|
| 1 | Anterior Thalamic Radiation (L) | **.0112*** | **628 (4.44%)** | .0631 |
| 2 | Corticospinal Tract (L) | **.0172*** | **630 (17.18%)** | .0631 |
| 3 | Superior Thalamic Radiation (L) | **.0172*** | **583 (13.51%)** | .0631 |
| 4 | Corticospinal Tract (R) | **.0368*** | **22 (0.57%)** | .0829 |
| 5 | Superior Longitudinal Fasciculus III (L) | **.0404*** | **30 (0.41%)** | .0829 |
| 6 | Arcuate Fasciculus (L) | **.0452*** | **2 (0.01%)** | .0829 |
| 7 | Corpus Callosum | .0922 | 0 (0%) | .1305 |
| 8 | Superior Thalamic Radiation (R) | .0982 | 0 (0%) | .1305 |
| 9 | Arcuate Fasciculus (R) | .1068 | 0 (0%) | .1305 |
| 10 | Superior Longitudinal Fasciculus III (R) | .1998 | 0 (0%) | .2198 |
| 11 | Anterior Thalamic Radiation (R) | .2294 | 0 (0%) | .2294 |

**Note.** p values are fixel-wise FWE-corrected within each tract. The minimum FWE-corrected p value from each of the 11 primary tract analyses was subjected to Benjamini–Hochberg FDR correction across tracts. qBH-FDR denotes the BH-FDR-adjusted tract-level p value. Bold values indicate $p < .05$ for within-tract FWE results and $q < .05$ for across-tract FDR-corrected results.

No tract reaches within-tract FWE significance or survives BH-FDR.

**Supplemental Table 5. Tract-of-Interest (TOI) Fixel-wise analysis of Group × Age interaction (CWS > controls).**

| Rank | Tract | Min. FWE-corrected *p* | FWE-significant Fixels, n (%) | $q_{BH\_FDR}$ |
|---|---|---|---|---|
| 1 | Anterior Thalamic Radiation (L) | **.0198***| **211 (1.49%)** | .2178 |
| 2 | Superior Thalamic Radiation (L) | .1410 | 0 (0%) | .7678 |
| 3 | Anterior Thalamic Radiation (R) | .2690 | 0 (0%) | .7678 |
| 4 | Superior Thalamic Radiation (R) | .2792 | 0 (0%) | .7678 |
| 5 | Corticospinal Tract (L) | .3688 | 0 (0%) | .8114 |
| 6 | Corticospinal Tract (R) | .7956 | 0 (0%) | .9842 |
| 7 | Superior Longitudinal Fasciculus III (L) | .9122 | 0 (0%) | .9842 |
| 8 | Arcuate Fasciculus (L) | .9332 | 0 (0%) | .9842 |
| 9 | Arcuate Fasciculus (R) | .9558 | 0 (0%) | .9842 |
| 10 | Corpus Callosum | .9678 | 0 (0%) | .9842 |
| 11 | Superior Longitudinal Fasciculus III (R) | .9842 | 0 (0%) | .9842 |
| **Note.** p values are fixel-wise FWE-corrected within each tract. The minimum FWE-corrected p value from each of the 11 primary tract analyses was subjected to Benjamini–Hochberg FDR correction across tracts. qBH-FDR denotes the BH-FDR-adjusted tract-level p value. Bold values indicate p < .05 for within-tract FWE results and q < .05 for across-tract FDR-corrected results. | | | | |
| No tract reaches within-tract FWE significance or survives BH-FDR. | | | | |

### Supplemental Figures

**Supplemental Figure 1: Fixel-Based Analysis Pipeline**. **A.** Diffusion-weighted MRI images underwent denoising and correction for *eddy* currents, motion, distortions, and bias field in native space. Fiber orientation distributions (FODs) were estimated using single-shell, multi-tissue constrained spherical deconvolution, and fiber density (FD) was calculated. **B.** A group FOD template was generated, and individual FOD images were nonlinearly registered to the template. Fiber-bundle cross-section (FC) was derived from the resulting transformations. A Fixel template and whole-brain tractogram were generated for connectivity-based Fixel enhancement. **C.** Fixel-wise statistical analyses were performed with family-wise error correction at $p < .05$. The acquisition parameters illustrated are for cohort 2; see Methods for parameters for both cohorts.

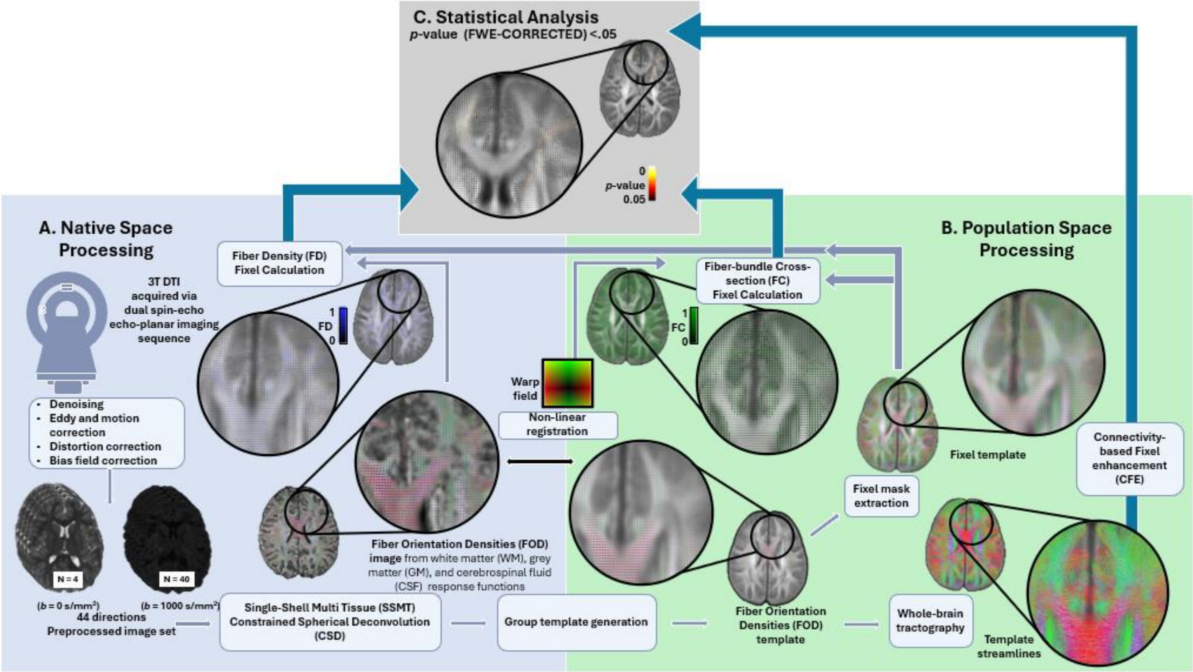

**Supplemental Figure 2. Group effects on Fixel-based measures in children who stutter (CWS) versus typically developing control participants. A.** (CWS > Controls) Axial, coronal, and sagittal slices show regions where children who stutter have *higher* logarithm-converted fiber cross-section (logFC) compared to controls; whole-brain significant Fixels are marked in red ($p < .001$) and yellow ($p < .005$). **B.** (CWS < Controls) Axial, coronal, and sagittal slices show regions where children who stutter have *lower* logarithm-converted fiber cross-section (logFC) compared to controls; whole-brain significant Fixels are marked in red ($p < .001$) and yellow ($p < .005$).

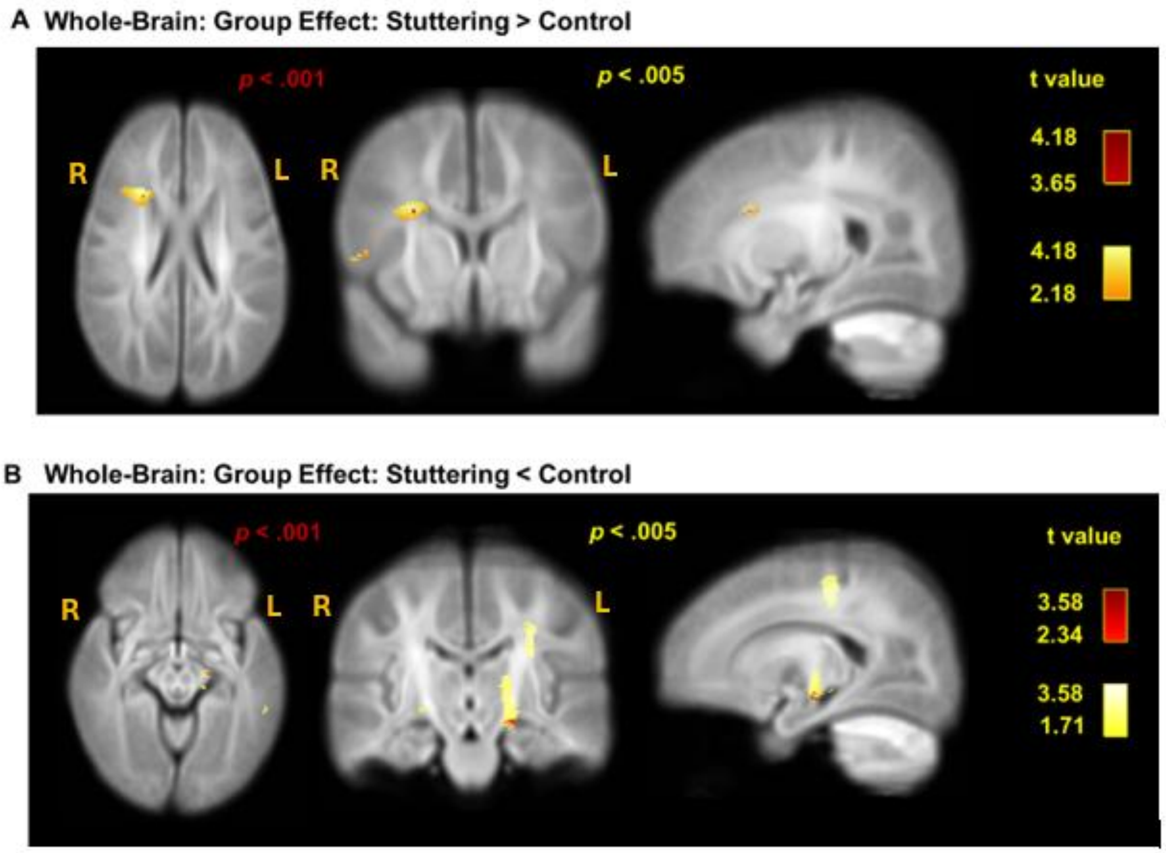

**Supplemental Figure 3. Reduced fiber cross-section in corticospinal and thalamocortical pathways in children who stutter.** Tract-of-interest analysis showing Fixels with lower fiber cross-section (FC) in children who stutter (CWS) relative to controls at the within-tract FWE-corrected threshold ($p_{FWE} < .05$; magenta), overlaid on the corresponding statistical maps. Effects were observed in the left arcuate fasciculus (AF), left anterior thalamic radiation (ATR), left superior longitudinal fasciculus (SLF) III, left superior thalamic radiation (STR), and bilateral corticospinal tracts (CST); however, none remained significant following BH-FDR correction across the 11 tract-of-interest analyses.

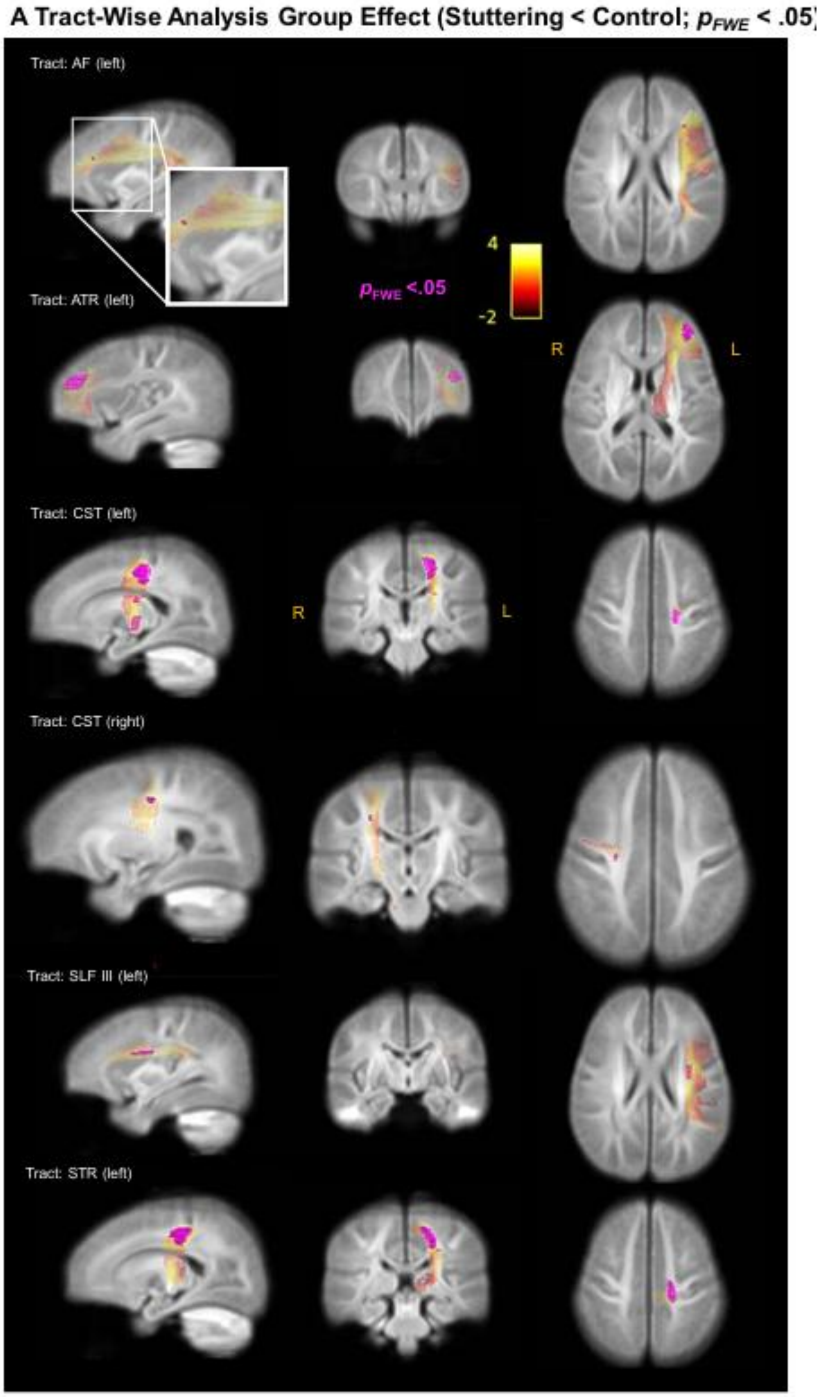

**Supplemental Figure 4. Group x Age Interaction effects on Fixel-based measures in children who stutter (CWS) versus typically developing participants. A.** (CWS > Controls) Axial, coronal, and sagittal slices show regions where children who stutter exhibit <u>*stronger*</u> age-related increase in logarithm-converted fiber cross-section (logFC) compared to controls. Whole-brain significant Fixels are marked in red ($p_{unc} < .001$) and yellow ($p_{unc} < .005$). **B.** (CWS < Controls) Axial, coronal, and sagittal slices show regions where children who stutter exhibit <u>*weaker*</u> age-related increase in logarithm-converted fiber cross-section (logFC) compared to controls. Whole-brain significant Fixels are marked in red ($p_{FWE} < .05$) and yellow ($p_{unc} < .001$).

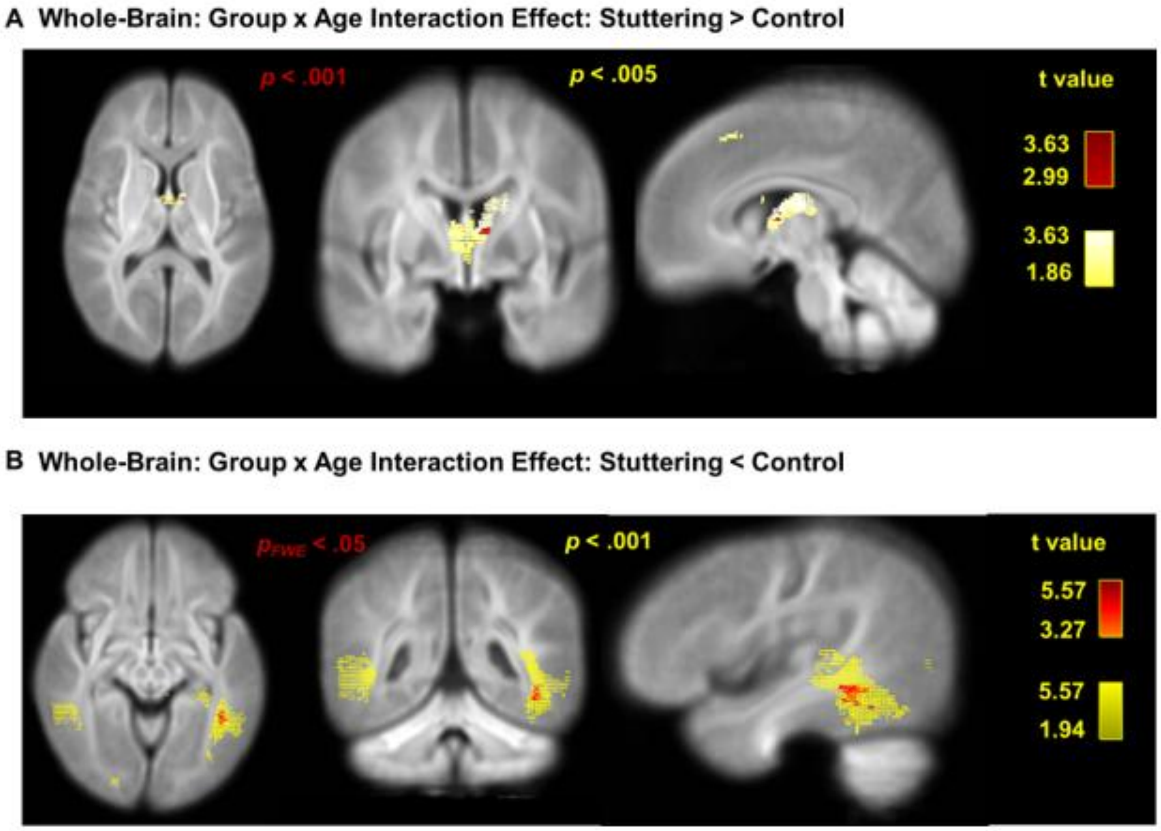

**Supplemental Figure 5. Tract-of-interest (TOI) Analysis: Group x Age Interaction effects on Fixel-based measures in children who stutter (CWS) versus typically developing control participants (CWS > Controls). A.** Tract-of-interest analysis showing Fixels with larger fiber cross-section (FC) in children who stutter (CWS) relative to controls at the within-tract FWE-corrected threshold ($p_{FWE} < .05$; magenta), overlaid on the corresponding statistical maps. **B.** Scatter plots showing relationship between Age (centered by group) and mean of significant Fixel logFC values for children who stutter for tract-of-interest analysis ($p_{FWE} < .05$). **C.** Scatter plots showing relationship between Stuttering Severity (SSI4) and mean of significant Fixel logFC values for children who stutter for tract-of-interest analysis. Abbreviations: **SSI4** = Stuttering Severity Instrument – version 4, **ATR** = Anterior Thalamic Radiation

**A Tract-Wise Analysis Group x Age Interaction (Stuttering > Control; $p_{FWE} < .05$)**

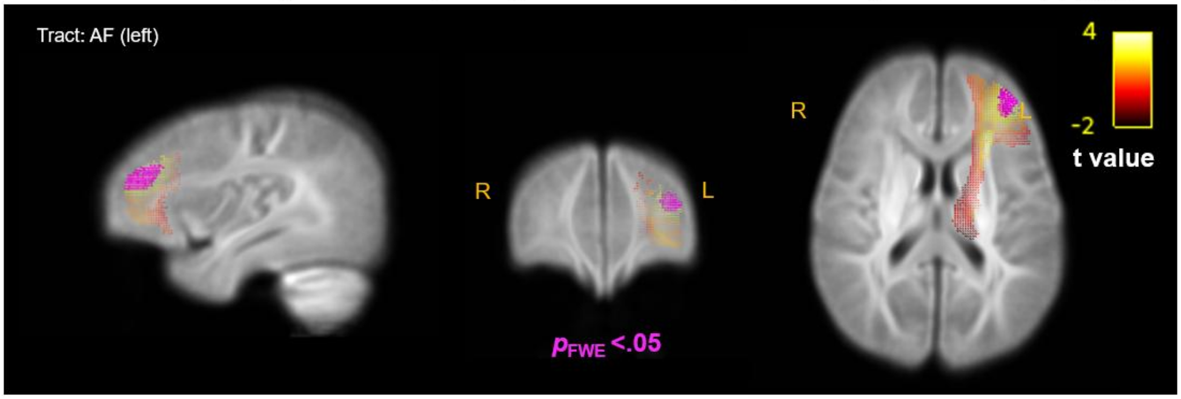


**B**

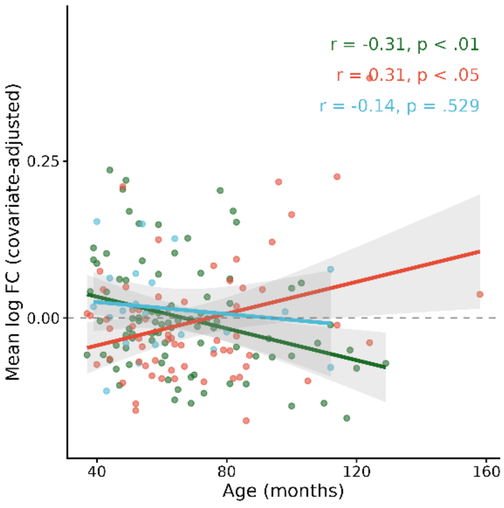


**C**

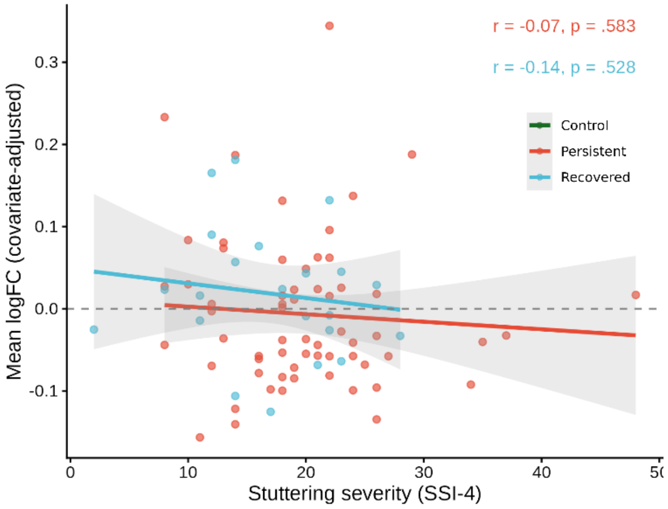

**Supplemental Figure 6. Tract-of-interest analysis of the Group × Age interaction in the right corticospinal tract. A.** Tract-of-interest analysis showing Fixels with a negative Group × Age interaction at the within-tract FWE-corrected threshold ($p$FWE < .05; magenta), overlaid on the corresponding statistical map. The right CST effect did not remain significant following BH-FDR correction across the 11 tract-of-interest analyses ($q$ = .1122). **B.** Age-related trajectories of covariate-adjusted log FC within significant Fixels in the left/right AF and right CST, illustrating the Group × Age interaction identified in the within-tract analysis. **AF**: arcuate Fasciculus; **CST**: corticospinal tract.

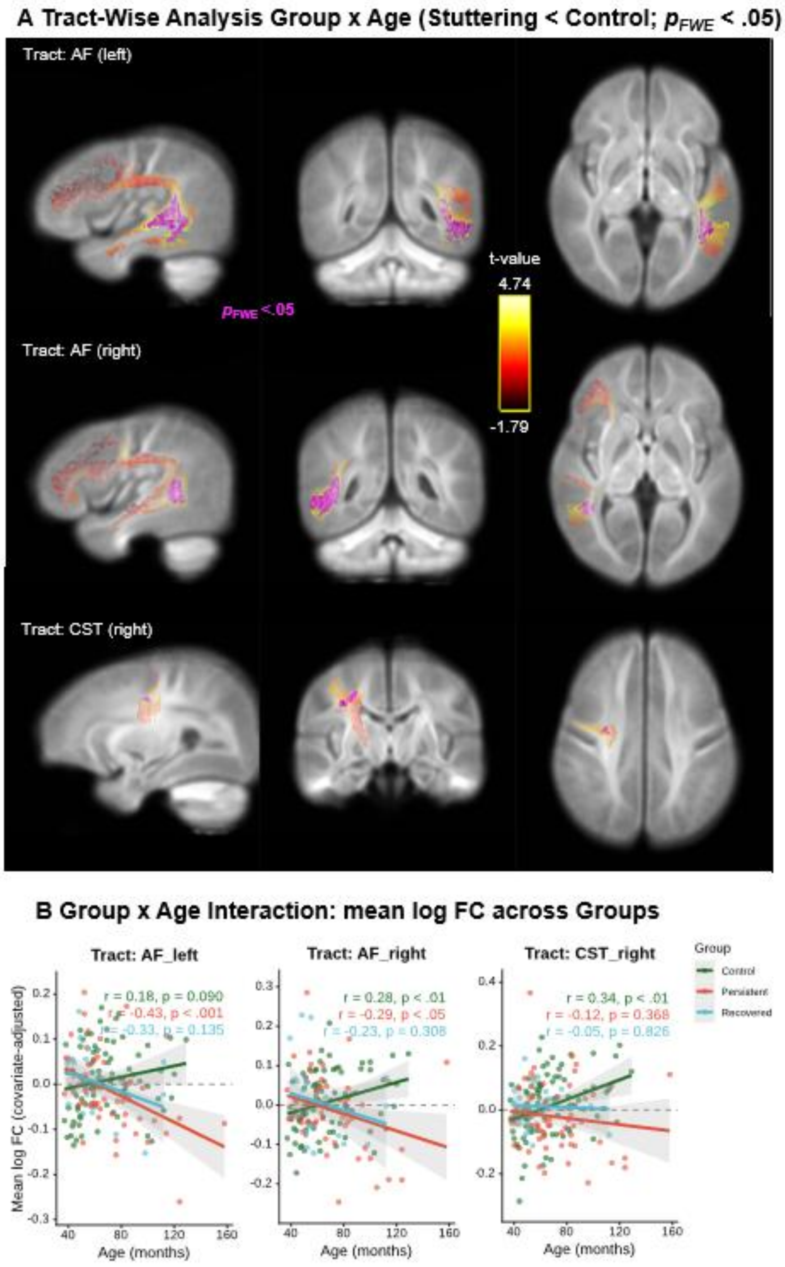
